# A complete RXFP1–relaxin interaction model unlocks the design of potent mini-protein modulators

**DOI:** 10.64898/2026.06.19.733483

**Authors:** Janik Clement, Tim Lkhagvajargal, Bradley L. Hoare, Tiffany Myint, Daniel R. Fox, Chunxiao Wang, Gavin J. Knott, Ross A.D. Bathgate, Rhys Grinter

## Abstract

Relaxin family peptide receptor 1 (RXFP1) is a multi-domain GPCR with compelling therapeutic potential, yet uncertainty surrounding the mechanism of its activation by the hormone H2 relaxin has hindered the development of selective modulators. Here, we combine deep learning based structural modelling with de novo protein design to overcome this barrier. We generate a high-confidence structural model of the RXFP1–relaxin complex that is strongly supported by existing biochemical and functional evidence. This model reveals that relaxin engagement stabilises the RXFP1 extracellular linker, thereby triggering receptor activation. Guided by this model, we design mini-protein modulators that either block linker stabilisation or enforce it and induce an active receptor geometry. These molecules act as potent, selective RXFP1 antagonists or agonists, achieving low-nanomolar activity in both engineered and endogenously expressing cell lines despite adopting folds unrelated to relaxin. Together, these findings define the mechanistic basis of RXFP1 signalling, establish the first de novo agonists and antagonists of this receptor, and demonstrate how AI-enabled modelling and design can target structurally complex GPCRs previously inaccessible to structure-guided drug discovery.

## Introduction

Relaxin family peptide receptor 1 (RXFP1) is a multi-domain G protein-coupled receptor (GPCR) that mediates diverse physiological effects in response to the hormone human relaxin-2 (H2 relaxin), including vasodilation, tissue remodelling and anti-fibrotic responses ^1–4^. RXFP1 belongs to the type C subfamily of leucine-rich repeat-containing GPCRs (LGRs), a subgroup of class A GPCRs. ^5^. It possesses a large relaxin-binding extracellular domain (ECD) comprising a leucine-rich repeat (LRR) domain and a low-density lipoprotein class-A (LDLa) domain, connected by a flexible interdomain linker, with each component critical for ligand recognition and signal transduction ^6^.

H2 relaxin is produced by the corpus luteum and placenta and is essential for maintaining pregnancy and parturition via RXFP1 ^7,8^. Beyond reproductive physiology, H2 relaxin regulates collagen turnover, promotes wound healing, and supports cardiovascular protection. These diverse biological roles have established RXFP1 as a promising therapeutic target for cardiovascular and fibrotic diseases ^9–13^. Agonists of RXFP1 show promise for treating heart failure, pulmonary and hepatic fibrosis, and other conditions where enhanced vasodilation or matrix remodelling is beneficial ^10,12,14–16^. Conversely, RXFP1 antagonists hold potential for disorders where aberrant tissue remodelling or tumour-associated stromal responses drive pathology, positioning RXFP1 as an emerging target in oncology and fibrotic disease biology ^1,16–18^. However, the development of relaxin-based therapeutics has been limited by inconsistent efficacy, underscoring the need for alternative strategies to modulate RXFP1 signalling ^11,19–21^.

A key barrier to drug development has been the limited structural information available for RXFP1. A cryo-EM structure of RXFP1 in its active-state has been determined, in which the transmembrane core is well defined, but the ECD and its interaction with relaxin is poorly resolved due to conformational heterogeneity ^22^. This limited understanding of relaxin engagement and RXFP1 activation has restricted the structure-guided design of modulators.

Advances in deep-learning-based structural modelling have enabled fast and accurate sequence-to-structure prediction of many complex protein–protein interactions, providing a powerful complement to experimental structural biology for drug discovery ^23–28^. Leveraging these capabilities, we used AlphaFold2 (AF2) to generate a high-confidence RXFP1–H2 relaxin model that predicts domain arrangements and interface contacts consistent with published biochemical, functional and NMR data ^12,29,30^. We used this model in AI-driven de novo protein design workflows to create mini-protein binders that precisely modulate RXFP1 activity ^31–34^. Biochemical and functional characterisation of these binders demonstrates that they are highly stable and function as intended, acting as either agonists or antagonists with low to sub-nanomolar potency. The fidelity of these modulators to their design validates the RXFP1-relaxin structural model, providing new mechanistic insight into receptor activation and establishing lead molecules for the development of next-generation RXFP1-targeted therapeutics.

## Results and Discussion

### The RXFP1-relaxin structural model strongly agrees with experimental data

The recently determined cryo-EM structure of RXFP1 bound to an engineered relaxin (PDB 7TMW) revealed key features of the transmembrane domain that underlie receptor activation ^12,22^. However, flexibility in the ECD precluded determination of the molecular basis for relaxin binding and receptor activation. To overcome this limitation, we used AlphaFold2 multimer to generate a full-length structural model of RXFP1 bound to H2 relaxin and evaluated its accuracy using available experimental data (Supplementary Data S1) ^12,25,26,29,30,35^. The model places the disulphide-linked relaxin A (H2A) and B (H2B) chains in complex with the concave surface of the RXFP1 LRR domain, where they simultaneously engage the interdomain linker such that the LDLa module is held close to the C-terminal end of the LRR domain (Figure 1a).

**Figure 1:**
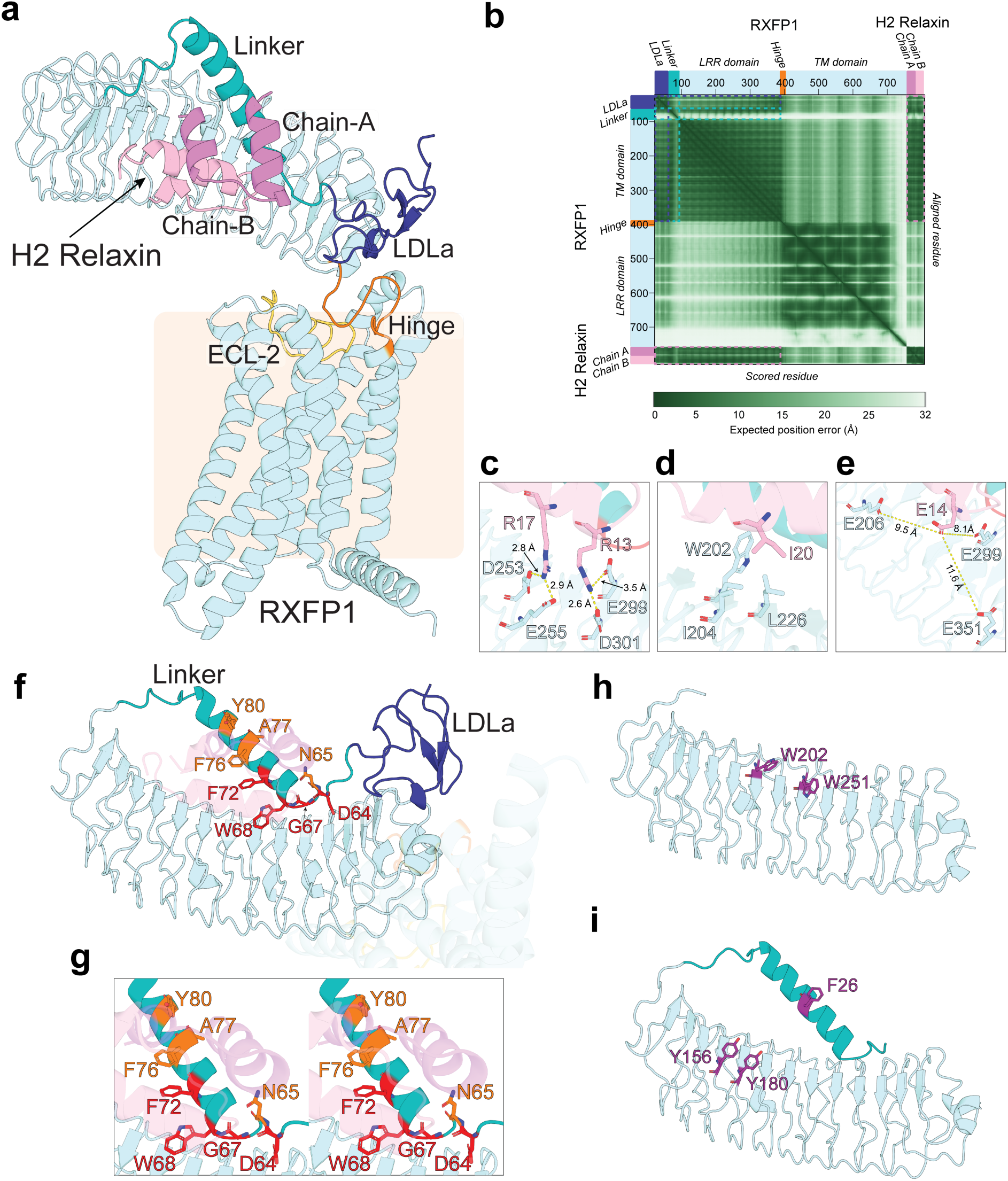
An AlphaFold2 model of the RXFP1-relaxin complex is a viable target for de novo binder design. a) An AlphaFold2 model of the RXFP1-H2 relaxin complex (ipTM= 0.93, pTM=0.76). The LRR domain (light blue) engages the relaxin B-chain (pink), and A-chain (magenta) with the interdomain linker (teal) connecting the LDLa module (dark blue). **b)** A Predicted Aligned Error (PAE) plot for the RXFP1-relaxin AlphaFold2 model showing the model-estimated positional. Key plot regions showing confidence of H2 relaxin (pink), linker (teal) and LDLa (dark blue) domain are highlighted. **c)** Charged interactions between R13 and R17 of relaxin chain B and negatively charged residues on the LRR of RXFP1. **d)** Interactions between I20 of relaxin chain B, and hydrophobic residues on the LRR of RXFP1. **e)** The proximity of E14 of relaxin chain B and glutamate residues on the LRR domain of RXFP1, which form crosslinks in the study by Erlandson et al. 2023 ^12^. **f)** A centred view of the RXFP1-ECD-H2 relaxin complex within the AlphaFold2 model. Residues highlighted in orange and red correspond to positions previously identified as important (orange) or essential (red) for RXFP1 activation by relaxin, based on analysis by Sethi et al. (2016 ^29^). These residues form a coordinated interaction network that positions the LDLa module for productive receptor activation. **g)** A cross-eye stereo view of the same interaction interface shown in panel f, illustrating how essential (red) and important (orange) linker residues pack against the LRR scaffold and H2 relaxin to stabilise the functional conformation of the linker. **h)** The target region of the RXFP1 model used for antagonist binder design. Highlighted residues (purple) represent the hotspot residues selected to define the antagonist target site on the LRR domain. **i)** The target region of the RXFP1 model used for agonist binder design, including the interdomain linker stabilised during RXFP1 activation. Residues highlighted in purple denote the hotspot residues chosen to guide design of agonist-mode binders intended to promote an activating linker configuration.

The model’s overall confidence is high, supported by an ipTM score of 0.93 and a global pTM score of 0.76. Per-residue confidence is uniformly high across the LRR domain and H2B (pLDDT >90) and remains strong for H2A (pLDDT >80; Supplementary Figure S1a). The relaxin–LRR interface is modelled with high confidence, as reflected by low inter-domain PAE values for both H2A and H2B relative to the LRR domain (5.53 Å and 5.12 Å, respectively; Figure 1b, Supplementary Figure S1b), indicating this is a reliable model of

The relaxin-engaged region of the extracellular linker is also confidently modelled (pLDDT >80), whereas the distal C-terminal segment (residues 77–92) shows reduced confidence (pLDDT 30–70), consistent with intrinsic flexibility. The spatial relationship between the linker and the LRR domain is nevertheless well defined (PAE 7.60 Å; Figure 1b, Supplementary Figure S1b). The LDLa module is predicted with high confidence overall (average pLDDT >80), although its alignment relative to the hinge region proposed to undergo LDLa-dependent rearrangement is supported with only moderate confidence (PAE 12.7 Å; Figure 1b, Supplementary Figure S1b) ^22^, indicating that mechanistic inferences in this region should be treated cautiously. Relaxin itself closely matches the experimentally determined crystal structure (PDB 6RLX), with an RMSD of 0.286 Å across 264 of 354 atoms and accurate modelling of all disulphide bonds, further supporting the reliability of the hormone–receptor complex ^36,37^.

The model relaxin-ECD complex strongly correlates with decades of experimental work. Relaxin B-chain residues R13 and R17 form electrostatic contacts with acidic residues E299/D301 and D253/E255 as predicted by early modelling (Figure 1c) ^35^. These interactions are consistent with mutagenesis data showing that both acidic pairs are required for high-affinity relaxin binding and that disrupting the E299/D301 pair abolishes relaxin binding and cAMP signalling in full-length RXFP1 ^30,35^. Likewise, our model predicts that relaxin B-chain I20 sits within a hydrophobic pocket formed by RXFP1 residues W202, I204, and L226 (Figure 1d). Alanine substitutions at these positions cause near-complete loss of relaxin binding for W202A, strong loss for I204A, and a moderate reduction for L226A ^35^. Our model also places relaxin B-chain E14 within ∼12 Å of LRR residues E206, E299, and E351, consistent with crosslinking observed between E14 and these residues (Figure 1e) ^12^. These data combined with the confidence of the model provide independent corroboration of the orientation of relaxin bound to the LRR in our model.

The agreement between the model and functional data extends to interactions between relaxin and the interdomain linker. The functionally critical 63-GDNNGW-68 motif of the linker is positioned centrally within the relaxin-ECD interface, with W68 buried between relaxin and the LRRs, providing a structural rationale for the severe affinity and signalling deficits observed in the RXFP1 W68A mutant (Figure 1f,g) ^5^. In addition, linker residues D64, G67, and F72 form direct stabilising interactions with relaxin or anchor the linker against the LRR domain, in strong agreement with the >10-fold reduction in relaxin affinity of full-length RXFP1 in alanine substitution mutants of these residues ^29^. In addition, alanine substitution of N65, F76, A77, and Y80, which are also positioned at the relaxin-linker interface in the model, significantly reduces the affinity of RXFP1 for relaxin (Figure 1f,g) ^29^. NMR analysis of the isolated LDLa–linker fragment supports the structural features of our model. In the absence of relaxin, the linker beyond C62 is largely unstructured, with only a short helical turn observed (Q71–K74). Relaxin binding induces helix propagation extending from L70 to S78, covering the majority of the helical region that interacts with both relaxin and the LRR domain in our model ^29^.

Together, these data support an RXFP1 activation model in which relaxin binding stabilises the inter-domain linker against the LRR domain, predominantly through interaction with a defined helical element. This traps the linker between relaxin and the LRR, suggesting that the linker must first interact with the LRR before relaxin binding. Linker stabilisation tethers the LDLa module at the C-terminal region of the LRR domain, positioning it near the hinge preceding the 7TM bundle, which contains residues essential for receptor activation (Figure 1a) ^12^. The tethering of the LDLa module in this position likely drives the structural rearranging of the TMD that elicits receptor activation (Figure 1a). Through this mechanism, relaxin binding at the LRR domain likely initiates a conformational cascade that propagates across RXFP1 and activates signalling.

The strong agreement between the high-confidence AlphaFold2 model and experimental evidence supports its use as a template for rational, de novo design of mini-protein modulators targeting the RXFP1 ECD.

### De novo binders antagonise RXFP1 by blocking relaxin and linker binding

To develop RXFP1 antagonists with potential therapeutic utility in oncology and fibrotic disease ^10,14–16,22^, we de novo-designed mini-protein binders targeting the extracellular LRR domain. Guided by the structural model of the RXFP1–relaxin complex, we hypothesised that binders engaging the relaxin- and linker-binding surface of the LRR domain would competitively inhibit relaxin binding and antagonise receptor activation. A target structural model encompassing residues 91–318 of the RXFP1 LRR domain was constructed, and two target tryptophan residues (W202 and W251), located within the predicted relaxin- and linker-binding interface, were used to guide binder localisation during design (Figure 1h). De novo binder design campaigns were performed using either RFdiffusion coupled with ProteinMPNN or the BindCraft pipeline, enabling direct benchmarking of these computational approaches ^31–34^. Final designs were filtered to avoid the N303 glycosylation site ^38^. 96 RFdiffusion-derived and 48 BindCraft-derived designs meeting program-defined quality criteria were advanced to experimental evaluation.

All candidate mini-protein binders were expressed and purified in parallel, with high overall expression observed for both platforms (Figure S2). RXFP1 binding was assessed using two complementary assays. Direct binding to the RXFP1 ECD was measured by biolayer interferometry (BLI), measuring response during association (200 s) and dissociation (500 s) phases (Figure 2a,b,c). Competition of binders with europium-labelled H2 relaxin (Eu-H2 relaxin) for binding to RXFP1 was quantified using a cell-based assay in HEK293T cells overexpressing the receptor (Figure 2d,e). A substantial fraction of binders from both design pipelines effectively displaced H2 relaxin in the competition assay, indicating high affinity interaction with the relaxin binding surface area of the RXFP1 LRR domain (Figure 2d,e). Success rates derived from the relaxin competition assay (cutoff <40% residual Eu-H2 relaxin binding) indicated that 24 of 48 BindCraft binders (50.0%) were strong relaxin competitors, compared with 23 of 96 RFdiffusion-derived binders (24.0%). Despite the lower overall success rate for RFdiffusion, both pipelines yielded multiple binders capable of near-complete relaxin displacement. A correlation was observed between a positive BLI binding response and relaxin competition (Figure 2f,g), corroborating binder identification between the two methods and indicating that binders engage the RXFP1 ECD and directly inhibit relaxin binding.

**Figure 2:**
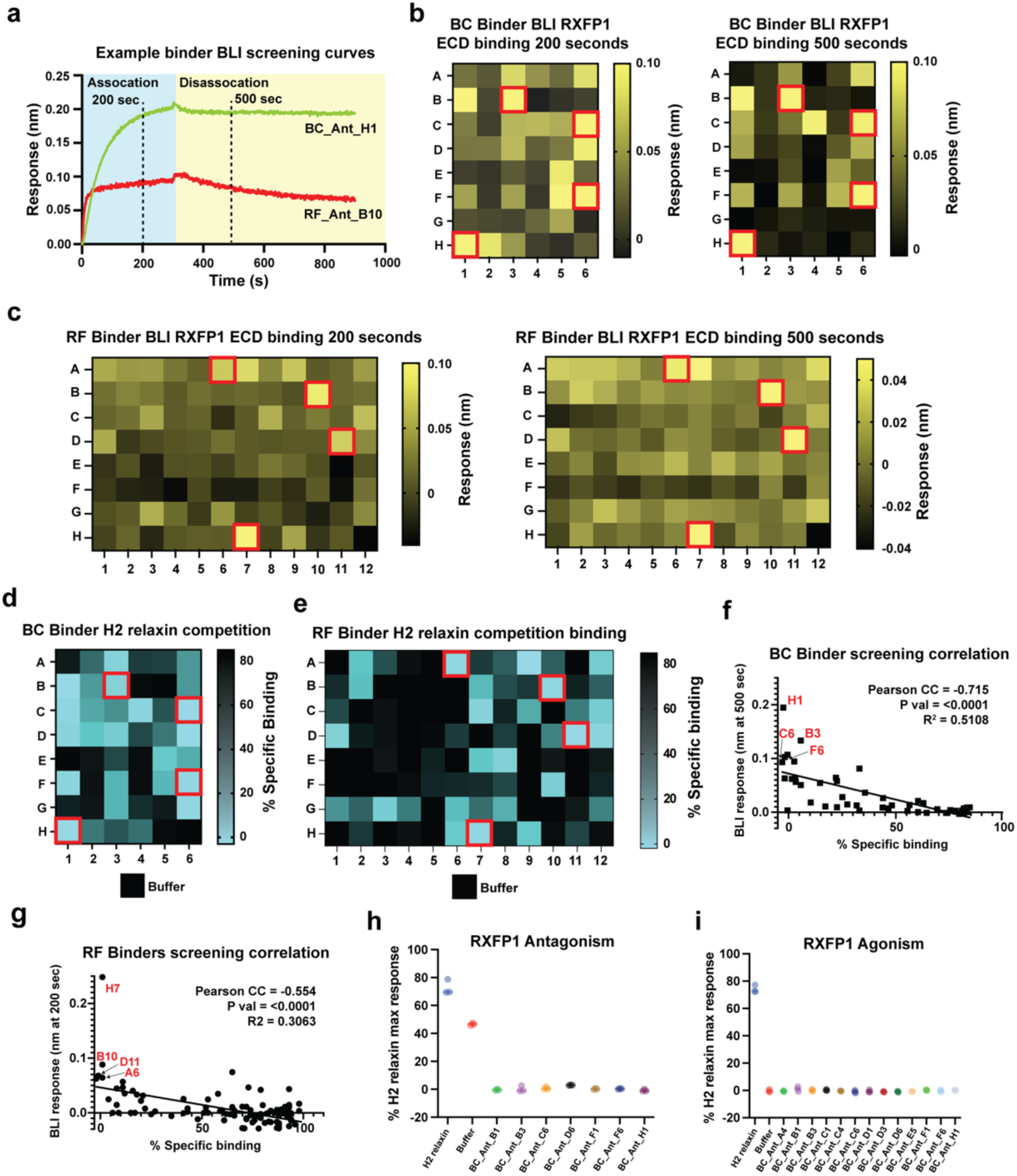
Screening of de novo RXFP1 antagonist candidates for receptor interaction. a) A BLI sensorgram plot showing response curves for two representative antagonist binders associating with the RXFP1 ECD. BLI Binding responses of the **b)** 48 BindCraft-designed antagonist binders and **c)** 96 RFdiffusion-designed antagonist binders 200 and 500 seconds after the start of the association step. Wells outlined in red designate binders with strong responses selected for further analysis. Competition between **d)** BindCraft binders and **e)** RFdiffusion binders, and europium-labelled H2 relaxin for binding to full-length RXFP1 in whole cells. The heatmap shows the percentage of specific competition relative to a buffer control. Correlation between BLI response at 500 seconds and percentage relaxin competition for **f)** the BindCraft and **g)** RFdiffusion binder sets. A negative correlation (BindCraft Pearson CC = −0.715, P < 0.0001, R² = 0.5108; RFdiffusion Pearson CC = −0.554, P < 0.0001, R² = 0.3063) indicates that stronger displacement of H2 relaxin corresponds to increased BLI binding response. **h)** Antagonist activity of selected BindCraft binders in a whole-cell assay measuring inhibition of H2 relaxin–induced RXFP1 signalling. Points show percent remaining relaxin response in the presence of each binder, with lower values indicating stronger antagonism. **i)** Agonist activity testing of the same binders reveals no detectable RXFP1 activation, consistent with their design as antagonists. 0.05 nM H2 relaxin was used in (h) and (i), corresponding to 80% of maximum RXFP1 activation in this system.

To determine whether high-affinity binders functionally antagonised RXFP1 signalling, a subset of the most potent BindCraft-derived candidates was evaluated in cell-based receptor activation assays. All tested binders suppressed H2 relaxin-induced RXFP1 signalling (Figure 2h), while none exhibited significant receptor activation in the absence of H2 relaxin (Figure 2i), indicating a lack of agonist activity and supporting a pure antagonistic mode of action.

Based on our primary screen, we selected the most effective mini-protein antagonists from each of the BindCraft and RFdiffusion design campaigns for detailed characterisation (Figure 3a,b). All eight candidates were expressed in *E. coli* and purified to homogeneity in milligram quantities and demonstrated high thermal stability and minimal aggregation across a 25-90 °C temperature range (Figure S3,S4).

**Figure 3:**
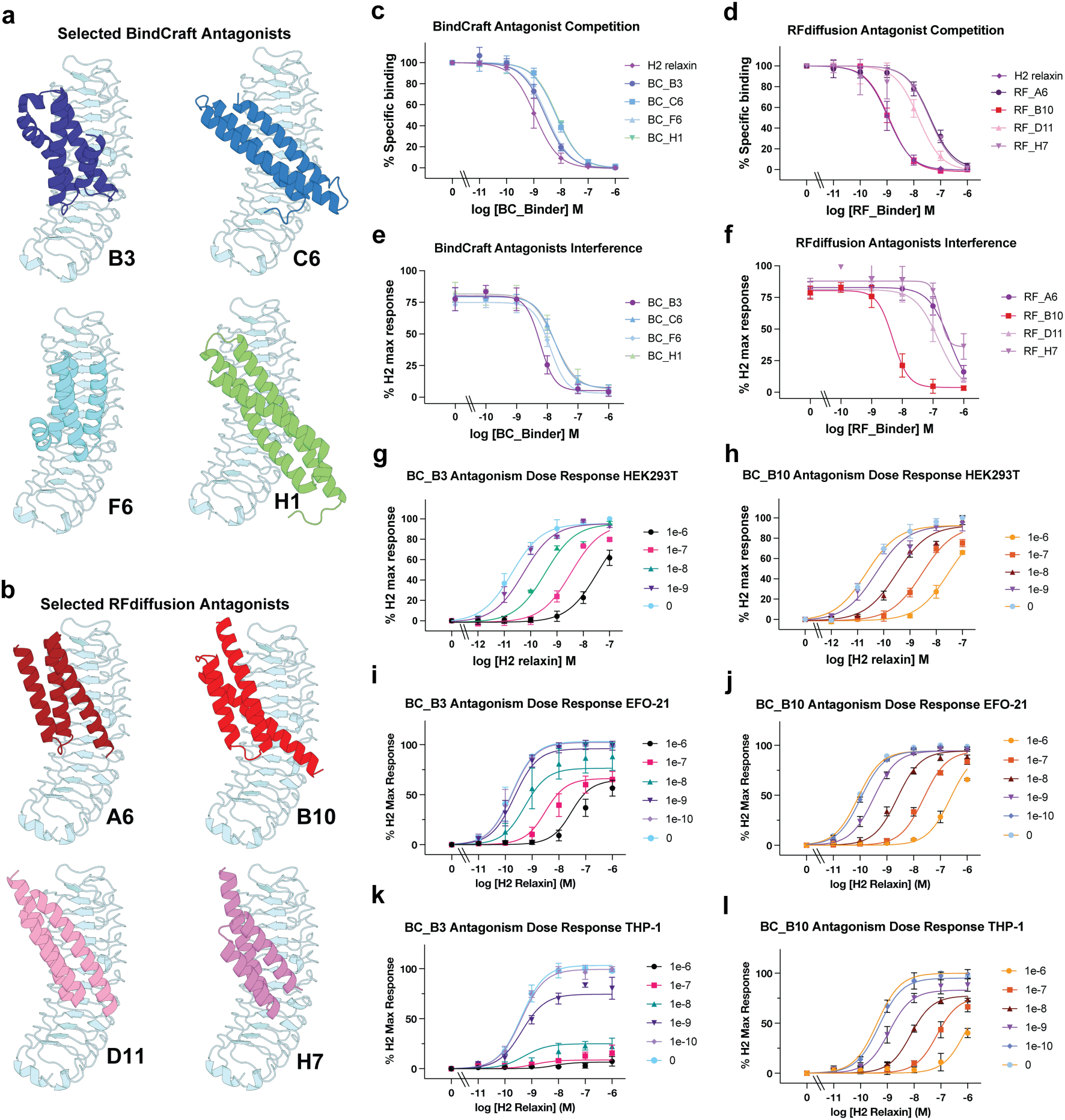
De novo–designed mini-protein antagonists bind the RXFP1 ECD and inhibit relaxin signalling. a,b) Structural models of top BindCraft and RFdiffusion-derived RXFP1 antagonist mini-proteins bound to the RXFP1 ECD target model. The RXFP1 ECD target is shown in light blue, individual antagonists coloured and labelled as indicated. Binders engage overlapping regions of the LRR domain corresponding to the predicted relaxin-binding surface. **c,d)** Competition binding assay titrations showing displacement of europium-labelled H2 relaxin from full-length RXFP1 in HEK293T cells by BindCraft and RFdiffusion-derived antagonists. Data are plotted as a percentage of specific H2 relaxin binding versus antagonist concentration. Unlabelled H2 relaxin competition is shown for comparison. **e,f)** Titrations of antagonism of RXFP1 signalling by BindCraft and RFdiffusion-derived antagonists measured using the HEK293T RXFP1 CRE reporter assay. **g,h)** Dose–response curves showing antagonism of H2 relaxin–induced cAMP signalling in RXFP1 expressing HEK293T cells by BC_Ant_B3 and RF_Ant_B10 measured using the HEK293T RXFP1 CRE reporter assay. **i,j)** Dose–response curves showing antagonism of H2 relaxin–induced cAMP signalling in EFO-21 cells by BC_Ant_B3 and RF_Ant_B10 measured using a real-time BRET-based cAMP sensor. **k,l)** Dose–response curves showing antagonism of H2 relaxin–induced cAMP signalling in THP-1 cells by BC_Ant_B3 and RF_Ant_B10. Increasing antagonist concentrations cause a rightward shift and reduction in maximal response, indicative of insurmountable antagonism in endogenous RXFP1-expressing cells. Data points represent mean ± S.E.M of replicate measurements from independent experiments. Quantitative parameters are reported in Table 1.

**Table 1:** Binding affinity and antagonistic potency of de novo–designed RXFP1 antagonist mini-proteins.

| *Binder* | *Competition Binding* |  | *Antagonism* |  |
| --- | --- | --- | --- | --- |
|  | *K<sub>i</sub> (nM)* | *95% CI of K<sub>i</sub> (nM)* | *IC<sub>50</sub>* | *95% CI of IC<sub>50</sub> (nM)* |
| *H2 Relaxin* | 0.42 | 0.33 to 0.54 | - | - |
| *BC_Ant_B3* | 0.998 | 0.76 to 1.30 | 5.90 | 3.34 to 10.41 |
| *BC_Ant_C6* | 2.52 | 1.99 to 3.15 | 17.88 | 13.16 to 24.29 |
| *BC_Ant_F6* | 1.15 | 0.91 to 1.45 | 12.45 | 9.15 to 16.93 |
| *BC_Ant_H1* | 2.72 | 2.32 to 3.19 | 17.37 | 9.88 to 30.54 |
| *RF_Ant_A6* | 15.22 | 9.93 to 22.9 | 357.10 | N/A* |
| *RF_Ant_B10* | 0.46 | 0.37 to 0.56 | 4.90 | 3.02 to 7.95 |
| *RF_Ant_D11* | 5.73 | 4.01 to 8.02 | 153.20 | 50.77 to 462.4 |
| *RF_Ant_H7* | 14.85 | 6.91 to 30.1 | 168.60 | N/A* |

\*CI is very large

We first quantified binding affinity using the Eu-H2 relaxin competition assay. Assuming single-site competitive binding, the measured inhibition constants (K_i_) directly reflect the equilibrium dissociation constants (K_D_) of the mini-protein antagonists. Binder affinity spanned nearly two orders of magnitude, with K_i_ values ranging from 0.46 to 15.22 nM. The three highest-affinity antagonists were RF_Ant_B10 (K_i_ = 0.46 nM), BC_Ant_F6 (K_i_ = 1.15 nM) and BC_Ant_B3 (K_i_ = 0.998 nM), which displayed competition curves close to that of unlabelled H2 relaxin. BC_Ant_C6 and BC_Ant_H1 showed intermediate affinity (K_i_ ≈ 2.52–2.72 nM), while RF_Ant_A6 and RF_Ant_H7 were weaker competitors (K_i_ ≈ 15 nM) (Figure 3c,d, Table 1).

We next assessed functional antagonism using a HEK293T cell-based CRE (HEK-RXFP1 CRE) reporter assay to measure inhibition of H2 relaxin-induced RXFP1 activation. Antagonistic potency correlated well with binding affinity. RF_Ant_B10 and BC_Ant_B3 were the most potent antagonists, with IC₅₀ values of 4.9 nM and 5.9 nM, respectively. BC_Ant_F6, BC_Ant_C6, and BC_Ant_H1 showed less potent antagonism (IC₅₀ ≈ 12–18 nM), whereas RF_Ant_D11, RF_Ant_H7, and RF_Ant_A6 were markedly less effective, with IC₅₀ values exceeding 150 nM (Figure 3e,f, Table 1). Notably, only the highest-affinity binders achieved near-complete suppression of signalling, indicating that tight engagement with the RXFP1 ECD is required for effective antagonism, consistent with the high affinity of its interaction with H2 relaxin ^29^.

None of the mini-protein antagonists activated RXFP1 in the HEK-RXFP1 CRE reporter assay in the absence of H2 relaxin (Figure S5a). Additionally, none of the antagonists affected forskolin-stimulated cAMP production, indicating that the observed inhibition is not due to non-specific suppression of adenylyl cyclase activity (Figure S5b). To further assess pathway specificity, we titrated the two lead antagonists, BC_Ant_B3 and RF_Ant_B10, in the presence of either forskolin or the small-molecule RXFP1 agonist AZD5462 (Figure S5c). No concentration of either antagonist altered RXFP1 activation by these compounds. As AZD5462 likely activates RXFP1 via direct binding to an allosteric site of the transmembrane domain ^22,39,40^, the lack of antagonistic effect is consistent with the mini-proteins acting selectively at the RXFP1 ECD. To evaluate the target specificity of BC_Ant_B3 and RF_Ant_B10, we tested their ability to antagonise insulin-like peptide 3 (INSL3) activation of RXFP2, which shares 60% amino acid sequence identity with RXFP1 ^7^. RXFP2 activation by 1 nM INSL3 was unaffected by either RXFP1 antagonist binder at 1 μM, indicating a high level of specificity for RXFP1 (Figure S5d).

We evaluated the antagonism of BC_Ant_B3 and RF_Ant_B10 in detail in the HEK-RXFP1 CRE reporter assay by performing concentration–response curve shift analysis. Increased concentrations of both antagonists produced progressive rightward shifts of the agonist curves, consistent with competitive inhibition (Figures 3g and 3h). Both BC_Ant_B3 and RF_Ant_B10 induced largely parallel shifts with minimal effect on maximal response, with Schild analysis yielding antagonist potency (pA₂) values of 8.91 (95% CI 8.76 –9.08) and 9.25 (95% CI 9.13–9.36), respectively. These values indicate effective antagonism at nanomolar concentrations even in the presence of high levels of H2 relaxin.

We next assessed BC_Ant_B3 and RF_Ant_B10 antagonist activity in cell lines which endogenously express RXFP1. For a high-receptor-expression context, we used a cAMP BRET sensor (CAMYEL) in EFO-21 ovarian cancer cells, which overexpress RXFP1 endogenously ^41^. Real-time cAMP responses to H2 relaxin were measured following preincubation with purified antagonist proteins. In EFO-21 cells antagonism was insurmountable, increasing concentrations of H2 relaxin failed to fully overcome inhibition by either BC_Ant_B3 or RF_Ant_B10, as indicated by both, a concentration-dependent dextral shift, and a reduction in maximal response (Figure 3i,j). Following preincubation with 100 nM antagonist, the relaxin E_max_ (95% CI) was reduced to 65.5% (58.0–77.7%) for BC_Ant_B3 and 84.8% (81.2–88.7%) for RF_Ant_B10. pA_2_ values in this assay were determined using the insurmountable antagonist equation, based on equiactive dose-responses (DR) at concentrations below the EC_50_ to generate a regression plot (Figure S5e) ^42^. The calculated pA_2_ values of 8.67 (95% CI 8.43–8.95) for BC_Ant_B3 and 9.24 (95% CI 9.04–9.48) for RF_Ant_B10 were comparable to those calculated in the HEK-RXFP1 CRE reporter assay.

We next assessed antagonist activity in THP-1 cells, a monocytic leukaemia cell line that expresses RXFP1 at near-physiological levels of approximately 275 receptors per cell ^43,44^. In this low-receptor-reserve system, treatment with 10 nM BC_Ant_B3 or RF_Ant_B10 produced more pronounced insurmountable antagonism than observed in EFO-21 cells (Figure 3k,l). The relaxin E_max_ (95% CI) was reduced to 24.0% (20.3–32.6%) by BC_Ant_B3 and to 74.7% (72.38–77.08%) by RF_Ant_B10, consistent with reduced receptor reserve enhancing insurmountable antagonism. Estimated pA_2_ values (95% CI) were 8.62 (7.12–10.72) for BC_Ant_B3 and 9.39 (9.12–9.74) for RF_Ant_B10, confirming potent RXFP1 antagonism in this physiologically relevant cell context (Figure S5f).

To investigate whether a kinetic mechanism could rationalise the differences in the observed reduction of maximum relaxin response between BC_Ant_B3 and RF_Ant_B10 in our cell-based assays, we used biolayer interferometry (BLI) to determine the binding kinetics of BC_Ant_B3 and RF_Ant_B10 to the purified ECD of RXFP1. Though BC_Ant_B3 and RF_Ant_B10 showed similar affinity for the ECD, BC_Ant_B3 is estimated to have a three-fold slower off-rate than RF_Ant_B10, suggesting the binding kinetics play a role in the larger E_max_ reduction by BC_Ant_B3 (Figure S5g,h). The slow dissociation of BC_Ant_B3 is consistent with previously described insurmountable antagonists, where a prolonged receptor blockade reflects a degree of suppression against maximum agonist responses ^45–48^. Considering RXFP1 does not undergo significant ligand-induced desensitisation ^49^, these results demonstrate that our de novo generated binders are potent antagonists of H2 relaxin in near native cell lines.

### Structure-guided design of RXFP1 agonist mini-proteins

Designing agonist binders for RXFP1 presents a more challenging problem than antagonist development. Our structural model and available biochemical data indicate that relaxin activates RXFP1 by simultaneously engaging the LRR domain and tethering the interdomain linker, thereby stabilising the LDLa domain in a signalling-competent position (Figure 1) ^5,29,30^. We reasoned that de novo mini-protein agonists would need to recapitulate this dual engagement to trap RXFP1 in a relaxin-bound-like active state.

To test this hypothesis, we generated a target derived from our model of the RXFP1–relaxin complex, encompassing residues 63–382 of the ECD (Figure 1i). This target model includes the helical region of the interdomain linker in the relaxin-bound conformation. Binder placement was guided using three hotspot residues positioned to enforce simultaneous interactions with both the regions, Y156 and Y180 within the LRR domain and F26 within the linker helix. Using this target model, we performed a BindCraft design campaign and selected the top 48 designs for small-scale expression and screening. As with the antagonist binders, agonist binder expression was high, with >80% of designs produced successfully.

Initial screening was performed using the Eu-H2 relaxin competition assay (Figure 4a) and RXFP1 agonism assays using the HEK-RXFP1 CRE reporter assay (Figure 4b). Eight binders exhibited potent competition with Eu-H2 relaxin, while five significantly activated RXFP1 signalling. The success rate was substantially lower than for antagonist discovery, likely reflecting the more stringent requirement for productive ECD conformational trapping. Notably, several binders efficiently competed with relaxin but failed to activate RXFP1 (e.g. BC_Ag_A4), suggesting they do not bind in a way that activates the receptor (Figure 4a,b). Despite this, competition binding and RXFP1 activation showed a strong correlation across the full screening dataset (Figure 4c). Four binders that activate RXFP1, and the non-activating binder BC_Ag_A4, were selected for detailed characterisation and produced in milligram quantities (Figure 4d). All five mini-protein binders demonstrated high thermal stability and low aggregation propensity, except BC_Ag_A4 and BC_Ag_A5, which showed some aggregation over 65 °C (Figure S6,7).

**Figure 4:**
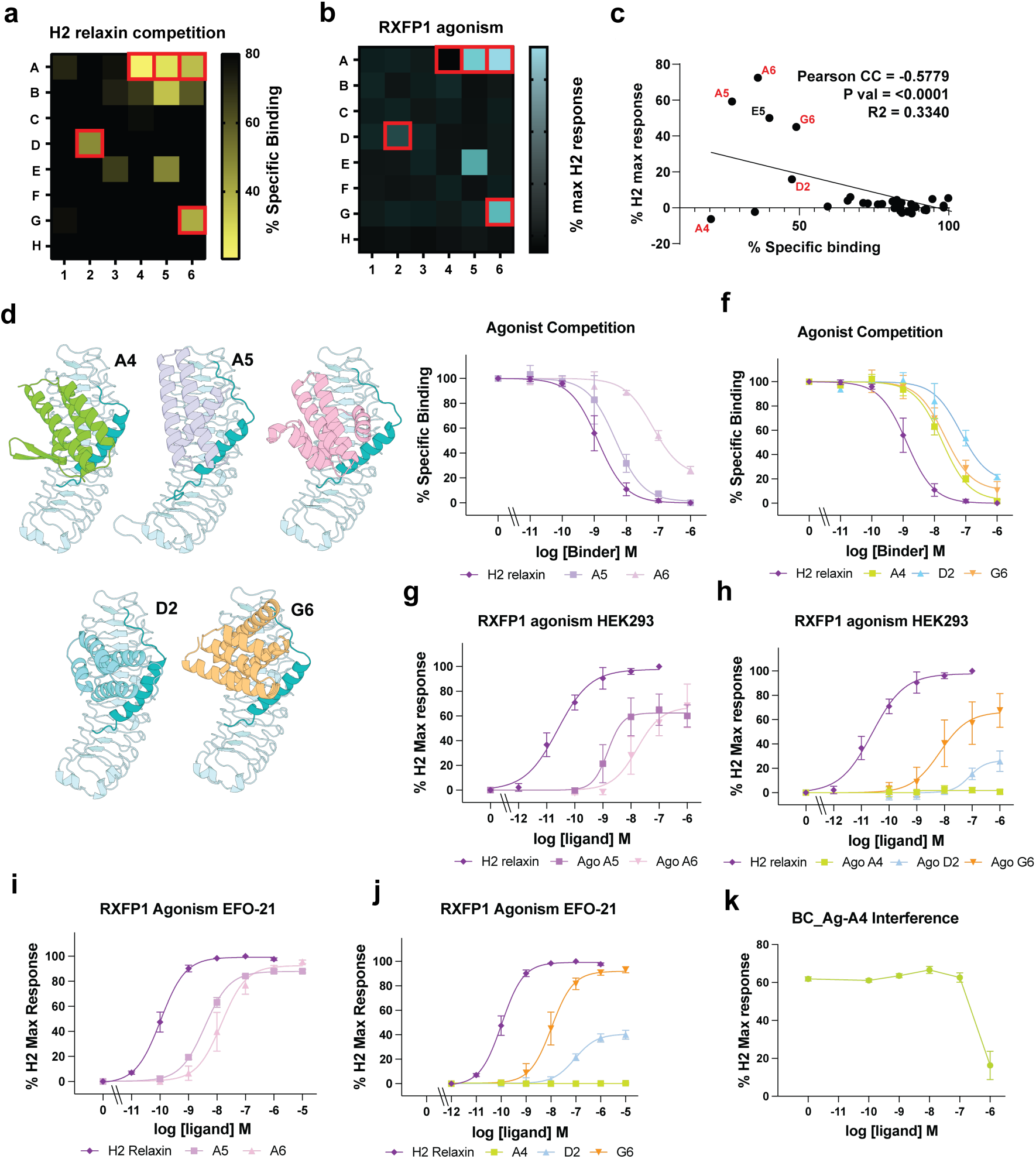
Functional characterisation of RXFP1 mini-protein agonist. a) A heatmap representation of competition binding to full-length RXFP1 in HEK293T cells, showing displacement of europium-labelled H2 relaxin by de novo–designed mini-protein agonists. **b)** A heatmap representation of RXFP1 agonism by mini-protein agonists in the HEK-RXFP1 CRE reporter assay. Binders selected for further characterisation are highlighted by red boxes. **c)** Correlation between RXFP1 agonist activity, measured as maximal cAMP response, and specific relaxin competition binding. Agonist binders (BC_Ag_A5, BC_Ag_A6, BC_Ag_G6 and BC_Ag_D2) cluster toward higher functional activity, whereas BC_Ag_A4 shows minimal activation despite substantial relaxin competition, indicating binding without productive receptor activation. **d)** Structural models of the selected ‘agonist’ binders bound to the RXFP1 ECD target region. RXFP1 is shown in light blue, with the interdomain linker highlighted in teal. Individual binders are coloured and labelled as indicated. **e,f)** Competition binding curves showing displacement of europium-labelled H2 relaxin by selected agonist binders. **g,h)** RXFP1 activation by selected agonist binders measured using the HEK-RXFP1 CRE reporter assay. Agonist binders BC_Ag_A5 and BC_Ag_A6 act as partial agonists relative to H2 relaxin, while BC_Ag_A4 shows no detectable activation. **i,j)** RXFP1 activation measured in EFO-21 cells using a real-time BRET-based cAMP sensor. In this endogenous RXFP1 context, BC_Ag_A5, BC_Ag_A6 and BC_Ag_G6 achieve full agonism relative to H2 relaxin, whereas BC_Ag_D2 shows limited activation and BC_Ag_A4 remains inactive. **k)** Interference assay showing that BC_Ag_A4 does not activate RXFP1 but weakly inhibits relaxin-induced signalling at higher concentrations, consistent with antagonist-like behaviour. Data points represent mean ± **S.E.M.** of replicate measurements from independent experiments. Quantitative binding and pharmacological parameters are reported in Table 2.

**Table 2:** Binding affinity and agonistic potency of de novo–designed RXFP1 agonist mini-proteins.

| *Binder* | *Competition Binding* |  | *Agonism in HEK-RXFP1 CRE Cells* |  | *Agonism Competition Response in EFO-21 Cells* |  |
| --- | --- | --- | --- | --- | --- | --- |
|  | *K<sub>i</sub> (nM)* | *95% CI of K<sub>i</sub> (nM)* | *EC<sub>50</sub> (nM)* | *95% CI of EC<sub>50</sub> (nM)* | *EC<sub>50</sub> (nM)* | *95% CI of EC<sub>50</sub> (nM)* |
| *H2 relaxin* | 0.49 | 0.39 to 0.61 | 0.025 | 0.018 to 0.035 | 0.11 | 0.094 to 0.126 |
| *BC_Ag_A4* | 6.58 | 5.08 to 8.64 | - | - | - | - |
| *BC_Ag_A5* | 1.74 | 1.11 to 2.65 | 1.49 | 0.84 to 3.59 | 3.79 | 3.37 to 4.26 |
| *BC_Ag_A6* | 25.21 | 15.71 to 38.91 | 16.62 | 6.35 to 119.7 | 15.80 | 10.45 to 24.95 |
| *BC_Ag_D2* | 25.8 | 13.46 to 45.89 | 79.50 | 29.37 to 271.6 | 87.88 | 64.49 to 125.1 |
| *BC_Ag_G6* | 79.8 | 48.80 to 132.7 | 6.82 | 1.92 to 53.15 | 10.77 | 7.73 to 15.4 |

Despite distinct amino acid sequences and folds, all selected binders engage RXFP1 in a way that recapitulates key features of relaxin binding. Each binder occupies a similar region of the LRR surface but does so using a unique structural solution. Notably, all agonists reproduce the important hydrophobic interaction of I20 of relaxin chain B, mediated by I98 (BC_Ag_A4), F15 (BC_Ag_A5), F21 (BC_Ag_A6), F54 (BC_Ag_D2), or L81 (BC_Ag_G6) (Figure S8a). In contrast, the electrostatic interactions formed by relaxin residues R13 and R17 with RXFP1 residues D253/E255 and E299/D301 are largely absent in the mini binder agonists, with only BC_Ag_D2 forming a comparable salt bridge via R52 to D253/E255 (Figure S8b). All agonists interact with the conserved aromatic residue W68, forming either direct hydrophobic contacts (BC_Ag_A4, BC_Ag_A5, BC_Ag_A6, BC_Ag_G6) or π-stacking interactions mediated by arginine and phenylalanine residues (BC_Ag_D2) (Figure S8c).

Competition binding titrations were performed on the selected binders with affinities spanning nearly two orders of magnitude (Figure 4e,f; Table 2). BC_Ag_A5 exhibited the highest affinity among the agonist series (K_i_ = 1.74 nM), whereas BC_Ag_A6 and BC_Ag_D2 bound with moderate affinity (K_i_ ≈ 25 nM). BC_Ag_G6 bound more weakly (K_i_ ≈ 80 nM), while BC_Ag_A4 exhibited sub-10 nM affinity despite its lack of agonist activity. Analysis of BC_Ag_A5 affinity for the isolated RXFP1 LRR domain by BLI indicated a K_D_ in the low nM range, consistent with competition binding data (Figure S9a).

Functional characterisation in the HEK-RXFP1 CRE reporter assay confirmed that four of the five binders acted as partial agonists. BC_Ag_A5 was the most potent, achieving an EC_50_ of 1.49 nM, followed by BC_Ag_G6 and BC_Ag_A6, which displayed EC_50_ values in the 7–17 nM range. BC_Ag_D2 was substantially weaker, with an EC_50_ of approximately 80 nM. Consistent with the screening data, BC_Ag_A4 did not produce detectable agonism in this assay (Figure 4g,h). To examine cell-type dependence, agonist activity was also evaluated in EFO-21 cells. In this context, BC_Ag_A5, BC_Ag_A6, and BC_Ag_G6 achieved full agonism relative to H2 relaxin, albeit with modestly higher EC_50_ values compared to HEK-RXFP1 cells (Figure 4i,j; Table 2), consistent with reduced receptor density. In contrast, BC_Ag_D2 showed only limited activation, and BC_Ag_A4 again failed to activate RXFP1. None of the mini-binder agonists activated the homologous relaxin-binding receptor RXFP2, at concentrations as high as 1 µM, indicating they are highly specific for RXFP1 (Figure S9b).

Interestingly, BC_Ag_A4 antagonised RXFP1 signalling at high concentrations (Figure 4k), with potency comparable to the weaker binders identified in the antagonist screen (Figure 3f). This behaviour is consistent with BC_Ag_A4 engaging the LRR domain at the relaxin binding site but failing to stabilise the interdomain linker in a signalling-competent configuration. Structural analysis does not provide a clear rationale for why BC_Ag_A4 may fail at linker stabilisation, as like H2 relaxin and the other agonist binders, it forms extensive interactions with the linker region (Figure S8d). However, it does indicate that subtle differences in engagement of the ECD and linker region, or steric constraints, can affect RXFP1 activation and switch functional outcomes between antagonism and agonism.

Overall, we have generated RXFP1-agonist mini-protein binders by explicitly targeting the relaxin-bound ECD conformation and enforcing simultaneous LRR and linker engagement, as predicted by our structural model. Although success rates are lower than for antagonist discovery, the identification of low-nanomolar agonists establishes a viable structure-guided strategy for engineering RXFP1 activators.

## Conclusions

In this study, we establish a high-confidence structural and mechanistic framework for RXFP1 activation and leverage it to design potent, de novo mini-protein modulators with defined agonist and antagonist activities. By integrating AlphaFold2-based modelling with deep-learning binder design, we show that stabilisation of the RXFP1 interdomain linker against the LRR domain by relaxin is the key structural determinant for receptor activation. Guided by this principle, we generate RXFP1 antagonists, which potently and selectively block relaxin signalling in both engineered and endogenously expressing cells, as well as de novo agonists that recapitulate relaxin-like activation despite adopting distinct folds and topologies. These modulators validate the RXFP1–relaxin structural model, providing direct experimental support for a linker-tethering mechanism of RXFP1 activation. This work illustrates how AI-driven structural modelling and protein design can overcome longstanding barriers in the development of modulators targeting multi-domain GPCRs, enabling the rational development of precision biologics for targets that have remained challenging for conventional drug discovery. More broadly, this work demonstrates that experimental structural information is not necessarily a prerequisite for structure-guided biologic discovery. The approaches and mechanistic insights presented here establish a generalisable paradigm for designing functionally selective protein modulators of complex receptors. Finally, this work has produced promising lead molecules for development as RXFP1-targeting therapeutics, expanding the therapeutic landscape for cardiovascular, fibrotic, and oncological diseases.

## Materials and Methods

### AlphaFold2 modelling

A structural model of the RXFP1-Relaxin complex was generated using Alphafold v2.3.2 implemented on the University of Melbourne Spartan Computing cluster. The Uniprot amino acid sequence entry for human RXFP1 (Q9HBX9) was used, as well as the amino acid sequence for mature H2 relaxin chains A and B derived from the structure of H2 relaxin (PDB ID = 6RLX). Five ranked models were produced by AlphaFold2, with relaxation performed on the top-ranked model. Models were compared for consistency, with all models converging on the same ECD-relaxin complex, with an average RMSD of 0.156 Å. The top-ranked model was used for further analysis and figure generation using PyMOL (Schrödinger). This model was trimmed to generate target models for mini-protein agonist and antagonist design. All models are provided in Supplementary Data 1.

### Predicted Aligned Error

PAE values were calculated as the mean of all PAE(i,j) entries where residue i belongs to the scored domain and residue j belongs to the aligned domain, extracted from AlphaFold2 JSON outputs. Depiction of Heatmaps were modified from ‘PAE viewer’ by Uni Göttingen ^50^. The PAE heatmap matrix is presented asymmetrically to preserve directional information, as PAE(A|B) ≠ PAE(B|A).

### RXFP1 mini-protein binder design

*Target model*: For mini-protein binder design, AlphaFold2-derived structural models of the human RXFP1 ectodomain were used as targets. For agonist design, a construct encompassing residues 63–382 of the ECD region, including the linker region containing the functionally important GDNNGWSL motif, was used as a target for binder design. For antagonist design, the ECD was further truncated to residues 91–382, excluding the linker region, to drive binder engagement on the LRR domain alone, including the region normally occupied by relaxin and the linker.

### Antagonist designs

For antagonist design, the RXFP1 ectodomain structures were used as targets for both BindCraft v1.5.0 and RFdiffusion-based binder design workflows ^31,32^.

*BindCraft workffow -* designs were generated using the surface-exposed RXFP1 LRR residues W202 and W251 as interaction hotspots. Binder lengths were set between 65 and 120 amino acids, and default models (e.g. default_filters.json, default_4stage_multimer.json) and parameters were used ^32^. A total of 318 binder models were generated, which passed all BindCraft quality filters, of which 69 passed a glycosylation exclusion filter requiring all binder atoms to be at least 6.5 Å from the glycan conjugated to N335 (corresponding to N303 in the signal peptide subtracted RXFP1 numbering scheme). Glycan position was identified by density associated with N335 in the low-resolution CryoEM map of the recent RXFP1 structure (EMD-26004) ^22^. The filtered designs were assessed for structural diversity, including length, helix topology, and secondary structure composition, interface size, and compatibility with downstream affinity capture tagging. From this set, 48 binders were selected for gene synthesis and experimental testing.

*RFdiffusion workffow -* RFdiffusion-based antagonist designs were generated using RXFP1 residues W202 and W251 as interaction hotspots. Default model weights were used, with both denoiser noise scale and frame scale set to zero. A total of 3,000 backbone designs were generated and subsequently passed through a deep-learning binder design pipeline for sequence generation using ProteinMPNN, followed by quality assessment using AlphaFold2 with an initial-guess protocol ^31,34^. Designs were screened for in silico success using an AlphaFold2 Predicted Aligned Error interaction score cutoff of less than 10. Approximately 200 designs passed this initial filter and were subjected to five rounds of iterative recycling through the sequence–structure refinement pipeline, yielding a final pool of 495 designs that met the same pAE interaction threshold. Designs were filtered to exclude models with any binder atoms within 7 Å of the predicted glycan associated with N335. The remaining 378 designs were manually curated to maximise structural diversity, interface geometry, and compatibility with affinity capture tagging. From this curated set, 96 binders were selected for synthesis and experimental testing.

### Agonist designs

*BindCraft workflow* - For agonist design, the RXFP1 ectodomain target structure was provided to BindCraft v1.5.0, with F26 (linker), Y156 (LRR), and Y180 (LRR) specified as interaction hotspots. Binder length range was set between 40 and 120 amino acids. BindCraft generated 551 models that passed its internal quality filters. Designs were filtered to exclude models with any binder atoms within 5.5 Å of the predicted glycan associated with N335, leaving 101 designs for further evaluation. These models were assessed for structural diversity, including length, helix topology, and secondary structure composition, interface size, and compatibility with affinity capture tagging. From this set, 48 agonist binder designs were selected for gene synthesis and experimental testing.

### Mini-protein binder expression and purification

#### Batch binder expression for screening

Binder gene sequences were synthesised and cloned into either pET28a(+) (N-terminal 6×His tag) or pET29b(+) (C-terminal 6×His tag) expression vectors using *NdeI* and *XhoI* restriction sites (Twist Bioscience). The position of the His tag was selected on a per-binder basis, with either N- or C-terminal tagging chosen to minimise disruption of the RXFP1 binding interface and to avoid perturbation of intramolecular contacts contributing to binder stability. Lyophilised plasmid DNA was reconstituted in 30 µL nuclease-free water to a minimum concentration of 7.86 ng µL⁻¹. Initial small-scale expression was performed in parallel. Expression plasmids were transformed into chemically competent *Escherichia coli* C41 (DE3) cells by heat shock in 2 mL 96-well plates (0.5 µL plasmid DNA at 10–50 µg mL⁻¹ added to 10 µL competent cells). Following transformation, 1 mL of LB broth was added, and cells were recovered at 37 °C for 1 h with shaking at 400 rpm. Kanamycin was added to a final concentration of 50 µg mL⁻¹, and cultures were incubated overnight at 37 °C. Transformed *E. coli* C41 (DE3) cells were used to inoculate 4 mL Terrific Broth (Merck) supplemented with 50 µg mL⁻¹ kanamycin in a 24-well plate format. Cultures were grown for 18 h at 30 °C before harvesting by centrifugation. Cell pellets were resuspended and lysed using B-PER reagent (Thermo Fisher Scientific) according to the manufacturer’s instructions. Lysates were clarified by centrifugation, and supernatants were transferred to 2 mL 96-well plates. Ni–agarose resin slurry (50 µL) was added to each well, followed by incubation at 4 °C for 1.5 h with shaking at 200 rpm. The resin–lysate mixtures were transferred to 96-well filter plates (30–40 µm pore size), and unbound material was removed using a vacuum manifold. Resin was washed three times with 1.5 mL wash buffer (50 mM Tris, 200 mM NaCl, 20 mM imidazole, pH 8.0). Bound proteins were eluted by incubation with 200 µL elution buffer (50 mM Tris, 200 mM NaCl, 500 mM imidazole, pH 8.0) for 5 min and collected into fresh 500 µL 96-well plates. Expression levels and purity were assessed by SDS–PAGE, and relative yields were estimated based on Coomassie-stained band intensities.

### Large-scale binder expression

Large-scale production was performed for selected RFdiffusion antagonist binders (RF_Ant_A6, RF_Ant_B10, RF_Ant_D11, RF_Ant_H7), BindCraft antagonist binders (BC_Ant_B3, BC_Ant_C6, BC_Ant_F6, BC_Ant_H1), and BindCraft agonist binders (BC_Ag_A4, BC_Ag_A5, BC_Ag_A6, BC_Ag_D2, BC_Ag_G6). Expression plasmids (1 µL) were transformed into 50 µL chemically competent *E. coli* BL21 C41 (DE3) cells by heat shock. Cells were recovered in 1 mL LB broth at 37 °C with shaking for 1 h. A 100 µL aliquot was plated onto LB agar supplemented with 50 µg mL⁻¹ kanamycin and incubated overnight at 37 °C to assess transformation efficiency (>100 colonies). The remaining culture was used to inoculate a 50 mL overnight LB culture containing 50 µg mL⁻¹ kanamycin. Overnight cultures were used to inoculate 2 L Terrific Broth and grown at 37 °C to an OD₆₀₀ of 0.8–1.0. Protein expression was induced with 0.3 mM IPTG, and cultures were incubated for an additional 16 h at 25 °C. Cells were harvested by centrifugation at 6,000 rpm for 20 min at 4 °C and stored at −80 °C. Frozen cell pellets were resuspended in lysis buffer (50 mM Tris, 200 mM NaCl, 20 mM imidazole, pH 7.9) supplemented with 1 mM MgCl₂, 0.1 mg mL⁻¹ lysozyme, 0.05 mg mL⁻¹ DNase I, and one quarter of a cOmplete protease inhibitor tablet (Roche). Suspensions were incubated on ice for 30 min and lysed using an EmulsiFlex cell disruptor (two passes). Lysates were clarified by centrifugation at 20,000 × g for 10 min at 4 °C and loaded onto a HiTrap nickel affinity column (Cytiva) pre-equilibrated in binding buffer. Proteins were eluted using a linear imidazole gradient (up to 500 mM imidazole) on an ÄKTA Start system (Cytiva). Fractions were analysed by SDS–PAGE and Coomassie staining. Fractions containing purified binder were pooled and further purified by size-exclusion chromatography using a Superdex 75 26/600 column (Cytiva) equilibrated in SEC buffer (50 mM Tris, 200 mM NaCl, pH 7.9) on an ÄKTA Pure system (Cytiva). Final fractions were analysed by SDS–PAGE, pooled, concentrated using 3 or 10 kDa MWCO centrifugal filters, and stored at −80 °C. Concentration was estimated by absorption at 280 nm using the sequence-based absorption coefficient calculated using ProtParam ^51^. Final yields ranged from 1.4 to 75 mg per binder per litre of culture.

#### Protein aggregation and thermal unfolding analysis

Back-reflection aggregation detection

Aggregation of purified binders were analysed using a Prometheus Panta instrument (NanoTemper Technologies). Back-reflection light scattering was recorded to determine aggregation onset temperatures. Samples were prepared in elution buffer at concentrations of 2–4 mg mL⁻¹ and loaded into standard capillaries. For binders H1 (BindCraft), B3 (BindCraft), and D11 (RFdiffusion), samples were diluted to 0.5 mg mL⁻¹ and loaded into high-sensitivity capillaries to minimise aggregation artefacts. Capillaries were heated from 25 °C to 95 °C at a rate of 1 °C min⁻¹ with continuous fluorescence and scattering measurements. Thermal parameters were extracted from processed scattering curves using Prometheus analysis software.

### Circular dichroism

Circular dichroism (CD) spectra were recorded using an Applied Photophysics CD spectrometer. Proteins were diluted to 0.01 mg mL⁻¹ in water, and spectra were collected from 260 to 190 nm at 25 °C. Due to aggregation propensity, binders D11 (RFdiffusion) and H1 (BindCraft) were further diluted to 0.005 and 0.0025 mg mL⁻¹, respectively. Thermal denaturation experiments were performed by heating samples from 25 °C to 90 °C at a rate of 1 °C min⁻¹. CD spectra (260–190 nm) were recorded at 5 °C intervals. Data were smoothed using Chirascan software (v4.2.14, Applied Photophysics) and plotted using GraphPad Prism 10.

#### Cell lines

Human embryonic kidney 293T (HEK-293T) cells stably expressing human RXFP1 or RXFP2 (hereafter referred to as HEK-RXFP1 and HEK-RXFP2 cells, respectively) were used for binding and signalling assays ^52^. HEK-293T cells stably expressing human RXFP1 were used together with a pCRE–β-galactosidase reporter construct which has been described previously ^53^. The human monocytic leukaemia cell line THP-1 expressing RXFP1 and stably transduced with the Epac-based BRET cAMP sensor CAMYEL (YFP–Epac–Renilla luciferase) was generated as described previously ^54,55^. The human ovarian tumour cell line EFO-21, which endogenously expresses RXFP1 ^41^, was also stably transduced with CAMYEL using lentiviral transduction following the same protocol as for THP-1 cells ^54^.

#### Cell culture and transfection

HEK-293T cells were maintained in Dulbecco’s Modified Eagle Medium (DMEM; Sigma-Aldrich) supplemented with 10% fetal bovine serum (FBS), 1% L-glutamine, and 1% penicillin–streptomycin. THP-1 cells were cultured in RPMI-1640 medium (ATCC formulation), and EFO-21 cells were cultured in RPMI-1640 supplemented with 20% heat-inactivated FBS, 1% L-glutamine, and 1 mM sodium pyruvate. All cell lines were maintained at 37 °C in a humidified incubator with 5% CO₂.

For real-time BRET assays, HEK-RXFP2 cells were transiently transfected with pcDNA3.1-CAMYEN (YFP–Epac–NanoLuc)^56^ using Lipofectamine 2000 (Thermo Fisher Scientific) according to the manufacturer’s instructions. Cells were seeded into assay plates 24 h after transfection.

#### Europium-labelled H2 relaxin competition binding assays

Binding of mini-proteins to RXFP1 was assessed using whole-cell competition binding assays with europium-labelled H2 relaxin (Eu-H2 relaxin) in HEK-RXFP1 cells. Assays were performed in poly-L-lysine–coated 96-well plates (Merck) as described previously ^57^. Semi-purified protein fractions were screened using 1 µL of neat extract together with a 1:10 dilution of the same sample. Control wells contained the equivalent volume of elution buffer. Purified proteins were tested in full competition binding curves spanning concentrations from 0.01 nM to 1 µM. Non-specific binding was determined in the presence of 1 µM unlabelled H2 relaxin (Corthera). Data are presented as mean ± S.D. of the percentage of specific binding from duplicate or triplicate wells and were reproduced in at least three independent experiments. Competition binding curves were fitted using a one-site binding model in GraphPad Prism 10. Statistical comparisons of pKᵢ values were performed using one-way ANOVA with Tukey’s multiple-comparison test.

#### HEK-RXFP1 CRE reporter gene assays

Agonist and antagonist activities of mini-proteins were assessed using a CRE-β-galactosidase reporter assay in HEK-RXFP1 cells as described previously ^57^. Semi-purified protein fractions that demonstrated RXFP1 binding were tested at 1 µL of neat extract and at a 1:10 dilution, either alone to assess agonist activity or in the presence of an approximate EC₇₀ concentration of H2 relaxin (0.05 nM) to assess antagonism. Control wells contained equivalent volumes of elution buffer, and maximal receptor activation was determined using 10 nM H2 relaxin. Purified antagonist proteins were tested in full dose–response inhibition curves (0.01 nM to 1 µM) in the presence of 0.05 nM H2 relaxin. To assess specificity, antagonists were also tested against an EC₇₀ concentration of the allosteric RXFP1 agonist AZD5462 (Cayman Chemical) or 5 µM forskolin. Schild analyses were performed using dose–response curves generated in the presence of increasing concentrations of H2 relaxin. Purified agonist proteins were tested in full dose–response activation assays over the same concentration range. Cells were incubated at 37 °C for 6 h, after which media were aspirated, and cells were frozen at −80 °C overnight. β-Galactosidase activity was quantified from cell lysates as described previously ^57^. All experiments were performed in duplicate or triplicate wells across at least three independent biological replicates. Dose–response curves were fitted using four-parameter sigmoidal models in GraphPad Prism 10 to determine pA_2_, pIC₅₀ or pEC₅₀ values. Schild plots were fitted using the Gaddum/Schild model with no Schild slope constraints.

#### Real-time kinetic cAMP activity BRET Assays

Real-time cAMP signalling was measured using Epac-based BRET sensors. HEK-RXFP2 cells transiently expressing CAMYEN and EFO-21 or THP-1 cells stably expressing CAMYEL were used for kinetic BRET measurements ^56^. Adherent HEK-RXFP2 and EFO-21 cells were plated in 96-well CulturPlates (Revvity) at 50,000 cells per well in assay buffer consisting of phenol red–free DMEM supplemented with 25 mM HEPES. The next day, culture medium was aspirated and replaced with assay buffer containing coelenterazine-H (NanoLight Technology) at a final concentration of 5 µM. Non-adherent THP-1 cells were plated at 80,000 cells per well in the same assay buffer. After a 3 h incubation, coelenterazine-H was added to achieve the same final concentration. Ligands were prepared at 10× final concentrations in assay buffer. Following substrate addition, purified antagonist proteins were added to achieve final concentrations ranging from 0.1 nM to 1 µM, with control wells receiving vehicle only. Plates were incubated at 37 °C for 5 min prior to measurement. BRET signals were recorded at 37 °C using a PHERAstar FSX plate reader (BMG Labtech) equipped with BRET1 Plus filters (emission 535/30 nm and 450/80 nm). BRET measurements were acquired at 1-min intervals, with five baseline cycles recorded before ligand addition and 25 cycles recorded thereafter. Purified agonist proteins were tested in full dose–response activation assays (0.1 nM to 10 µM). For antagonism experiments, concentration–response curves for H2 relaxin (0.01 nM–1 µM) or INSL3 (0.001 nM–100 nM) were generated in the presence of fixed antagonist concentrations. Forskolin (10 µM) served as a positive control. BRET ratios (535/450) were multiplied by 10,000 and ligand induced BRET ratios were baselined against vehicle-treated controls. Data from individual plates were obtained in duplicate wells across three independent experiments. Dose–response curves were derived from area-under-the-curve values, normalized to maximal agonist response (100%) and vehicle (0%), and fitted using three-parameter non-linear regression. In the endogenously RXFP1-expressing cells, the depression of maximal H2 relaxin response induced by B3 and B10 antagonism was indicative of insurmountability/hemi-equilibrium conditions, therefore, the data was fit to the following operational model as described previously ^48^:

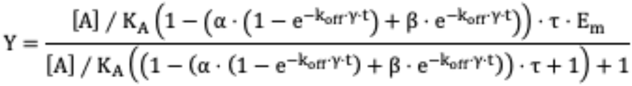

Where:

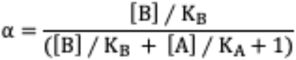

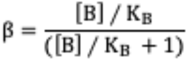

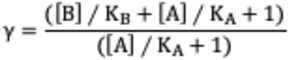

Where [A] and [B] represent the concentration of H2 relaxin and the antagonist (BC_B3 or RF_B10), respectively; K_A_ and K_B_ are the respective equilibrium dissociation constants; k_off_ denotes the rate of dissociation for the antagonist (min^-1^); t represents the assay incubation time (minutes), τ is the operational efficacy of H2 relaxin (comprising cell- and agonist-dependent properties), and E_m_ is the maximal system response. All parameters were shared across all data sets except t, which was fixed to the assay incubation of 25 minutes, and K_A_, which was set to the K_i_ value determined from the competition binding assay.

#### ALFA-tagged RXFP1 ectodomain production and purification

An expression construct for secreted RXFP1 ectodomain (ECD) protein was generated as a Geneblock dsDNA fragment flanked by BamHI/XhoI restriction sites and ligated into a lentiviral vector plasmid based on pCSC SP-PW (Addgene Accession #12337) which contained an IRES element to allow co-expression of mCherry fluorescent protein as an expression marker. The sequence coded for an IgGk cleaved signal peptide followed by an ALFA tag (SRLEEELRRRLTE) ^58^, residues 23-404 of RXFP1, and a C-terminal His_6_-tag. A stable HEK293F suspension cell line with constitutive secretion of this RXFP1 ECD protein was generated by lentiviral transduction of HEK293F suspension cells with subsequent FACS sorting to select for the mCherry positive population. Lentivirus was produced by co-transfection of HEK293T cells with the lentiviral packaging and envelope plasmids pMDL, pRSV-Rev, and pCMC-VSV-G as previously described ^54^. This stable cell line, named HEK293F-ectoRXFP1 was then maintained in suspension cell culture and passaged every 2-3 days as required.

For protein expression and purification, a fresh 500 mL culture of HEK293F-ectoRXFP1 cells at 2 million cells/mL was left to grow and secrete protein at 37 °C for 48 hours. The conditioned culture media was then harvested by centrifugation at 1000 x g for 10 minutes at 4 °C. The cell pellet was discarded and the supernatant further clarified by vacuum filtration through a Glass Whatman GF/C filter before addition of 20 mM HEPES (pH 7.4), 150 mM NaCl, and 2 mM CaCl_2_. The His-tagged protein was then purified with TALON® Metal Affinity Resin using a purification buffer consisting of 20 mM HEPES pH 7.4, 150 mM NaCl, 2 mM CaCl_2_ and 1% (w/v) glycerol. Protein was eluted with 200 mM imidazole and elution fractions containing protein were concentrated with Amicon 3kDa MWCO spin columns to a concentration of at least 0.5 mg/mL as determined by protein A280 measurement. Glycerol was then added to 10% (w/v) and aliquots of protein were flash frozen in liquid nitrogen and stored at –80 °C until required.

#### Biolayer interferometry analysis

Biolayer interferometry (BLI) experiments were performed using an Octet R8e system (Sartorius) with streptavidin biosensor tips. Sensors were saturated with 1–5 µM biotinylated anti-ALFA nanobody, expressed and purified as described previously ^59^, in binding buffer containing 20 mM HEPES pH 7.4, 200 mM NaCl, 0.1% (w/v) BSA, 0.02% (v/v) Tween-20, and 0.02% (w/v) NaN₃. All measurements were conducted at 25 °C with orbital shaking at 100 rpm during association and dissociation steps. ALFA-tagged RXFP1 ectodomain was diluted to 6 µg mL⁻¹ in binding buffer and immobilized onto biosensors to a loading response of 0.8–1.0 nm. Non-loaded sensors were used as reference controls. Following a 60-s baseline, sensors were exposed to mini-proteins for 300 s to measure association and then transferred to buffer-only wells for 600 s to measure dissociation. Data were reference-subtracted, baseline-corrected, inter-step corrected, and Savitzky–Golay filtered using Octet Analysis software. Binding curves were fitted using an association–dissociation model and plotted in GraphPad Prism 10.

## Data Availability

All data and structural models required to support the findings of the study are provided in this article or can be obtained from the authors on request.

## Acknowledgements

R.G. and D.R.F. are members of the Australian Research Council Industrial Transformation Training Centre for Cryo-Electron Microscopy of Membrane Proteins for Drug Discovery (IC200100052). R.G. was funded by an NHMRC EL1 investigator grant (APP1197376). G.J.K. was supported by the Snow Medical Research Foundation (SMRF2021-276). J.C. was supported by a University of Melbourne Scholarship. T.L. was supported by the Commonwealth through an Australian Government Research Training Program Scholarship. Work at the Florey Institute was supported by the Victorian Government Operational Infrastructure Support Program. The authors also acknowledge the use of the facilities at the Melbourne Protein Characterisation Facility, of the Bio21 Molecular Science C Biotechnology Institute, and the MBC Flow Cytometry Facility at the University of Melbourne and thank Dr Belinda Michell and Dr Troy Attard for training and assistance. The authors thank Sharon Layfield for technical assistance at the Florey Institute.

## Author Contributions

Conceived and designed the experiments: J.C., T.L., B.L.H., G.J.K., R.A.D.B, and R.G.; performed the experiments: J.C., T.L., B.L.H., D.R.F., C.W., and R.G.; analysed the data: J.C., T.L., B.L.H., D.R.F., R.A.D.B, and R.G.; contributed reagents/materials/analysis tools: R.A.D.B, G.J.K., and R.G.; wrote and edited the manuscript: J.C., T.L., R.A.D.B, and R.G.; acquired funding and provided project supervision: R.G., R.A.D.B., and G.J.K.; All authors edited and approved the manuscript.

## Supplemental Data

**Supplemental Data 1: AlphaFold2 model of the H2 relaxin-RXFP1 complex**

**Supplemental Data 2: RXFP1 mini-protein modulator sequences**

**Supplemental Data 3: Successful RXFP1 mini-protein modulator structural models**

**Supplemental Data 4: ALFA-tagged RXFP1-ECD sequence**

ORF sequence for ALFA-tagged RXFP1 ECD used in this study inserted into a lentiviral vector plasmid. Cleaved signal peptide; ALFA tag; RXFP1_23-404_; His_6_ tag

>METDTLLLWVLLLWVPGSTGPSRLEEELRRRLTEPGSQDVKCSLGYFPCGNITKCLPQLLHCN GVDDCGNQADEDNCGDNNGWSLQFDKYFASYYKMTSQYPFEAETPECLVGSVPVQCLCQG LELDCDETNLRAVPSVSSNVTAMSLQWNLIRKLPPDCFKNYHDLQKLYLQNNKITSISIYAFRGL NSLTKLYLSHNRITFLKPGVFEDLHRLEWLIIEDNHLSRISPPTFYGLNSLILLVLMNNVLTRLPDK PLCQHMPRLHWLDLEGNHIHNLRNLTFISCSNLTVLVMRKNKINHLNENTFAPLQKLDELDLG SNKIENLPPLIFKDLKELSQLNLSYNPIQKIQANQFDYLVKLKSLSLEGIEISNIQQRMFRPLMNL SHIYFKKFQYCGYAPHVRSCKPNTDGISSLENLLAGSHHHHHH*

## Supplemental Figures

**Figure S1:**
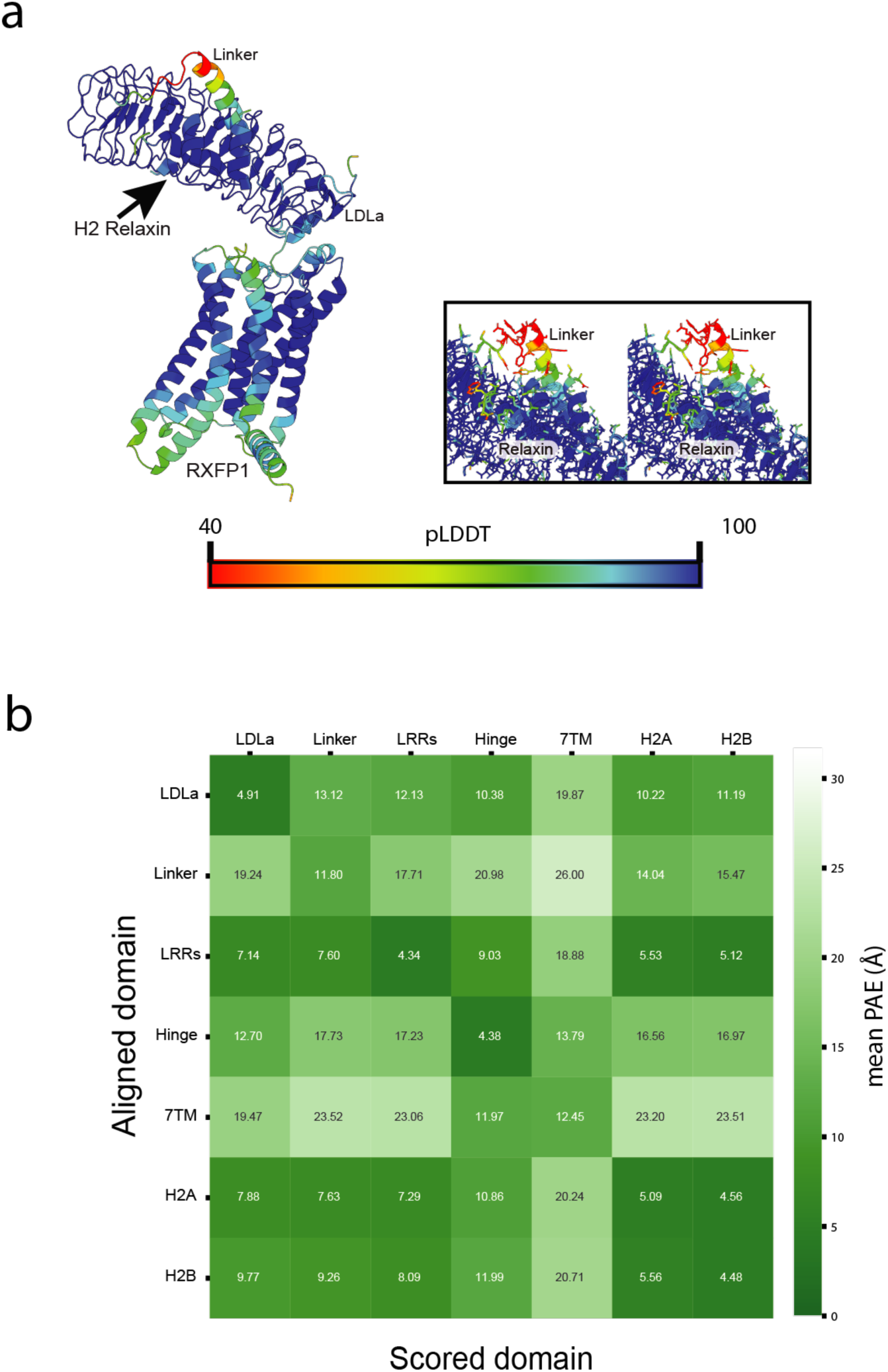
Predicted Confidence of the RXFP1-H2 Relaxin Complex Model. a) Per residue pLDDT confidence scores for the RXFP1–H2 relaxin complex AlphaFold2 Model. AlphaFold2-predicted model of full-length RXFP1 bound to H2 relaxin, coloured by per-residue predicted Local Distance Difference Test (pLDDT) scores (scale shown). High pLDDT values (>90) are observed across the RXFP1 leucine-rich repeat (LRR) domain and H2 relaxin, indicating high confidence in the predicted fold and interaction interface. The interdomain linker displays regionally variable pLDDT scores, with elevated confidence in the relaxin-bound helical segment and lower confidence in distal regions, consistent with conformational flexibility. The LDLa module is predicted with moderate to high confidence and is positioned proximal to the C-terminal end of the LRR domain. Insets show close-up views of the relaxin–ECD interface, highlighting the high-confidence regions that define the predicted ligand-binding and linker-stabilisation architecture. This confidence analysis supports the use of the model as a structurally reliable template for structure-guided binder design while appropriately delimiting flexible regions. **b)** Inter-domain resolved PAE Matrix for H2 relaxin bound RXFP1. Values represent the mean predicted aligned error (Å) for residues in the scored domain (column) when the structure is aligned on the aligned domain (rows). Lower PAE values (darker green) indicate higher confidence in the relative positioning of domain pairs. Diagonal blocks represent intra-domain confidence, while off-diagonal blocks represent inter-domain interface confidence. The scale bar indicates mean PAE (Å) from 0-31,7. AlphaFold models have been created based on UniProt ID: Q9HBX9. Artificial segmentation of RXFP1 following Erlandson et al. (2023). PAE matrix and mean PAE values established using the highest AlphaFold confidence model and PAE viewer ^50^.

**Figure S2:**
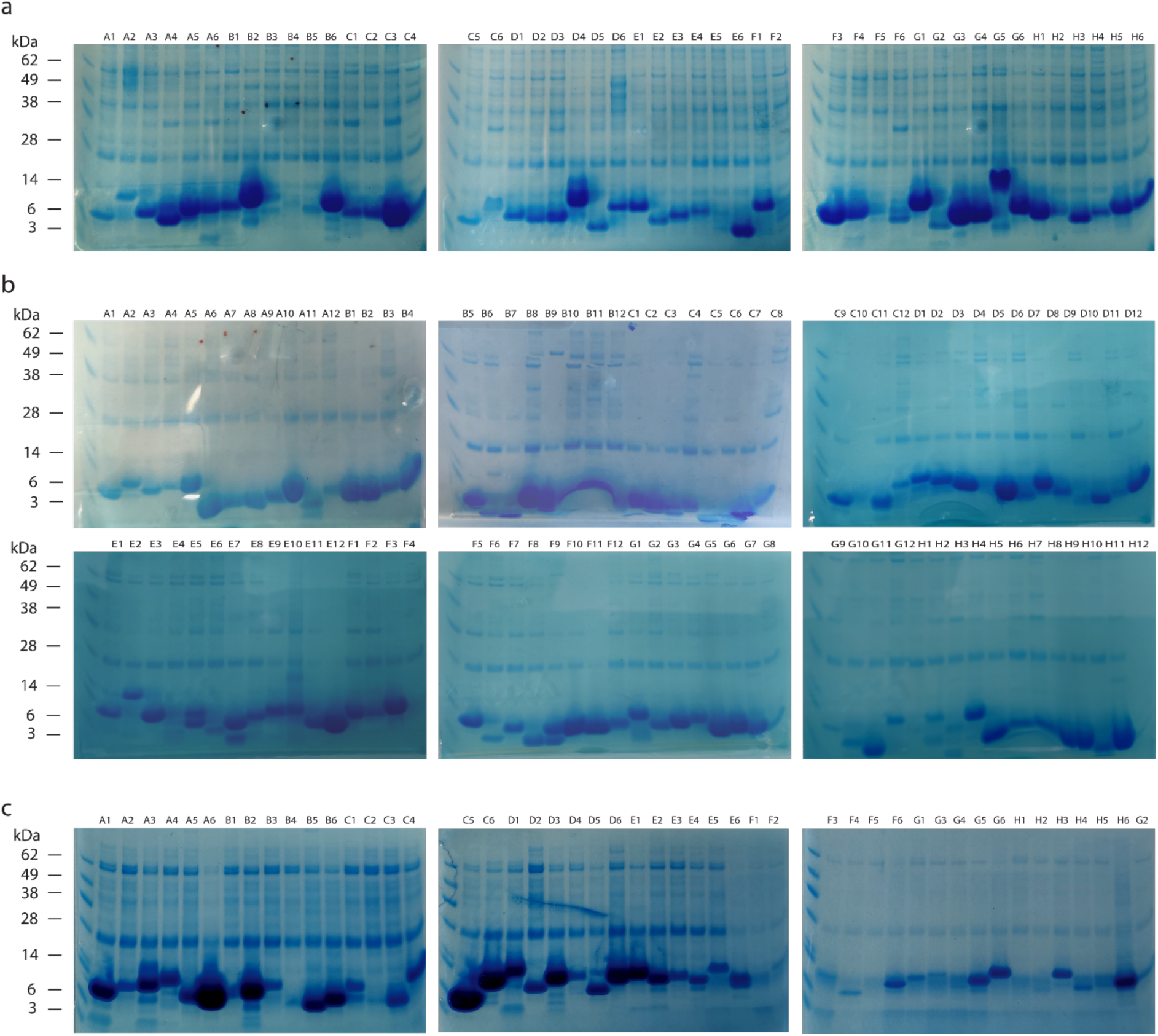
SDS-PAGE analysis of recombinant mini-protein batch expression in *E. coli*. SDS-PAGE results for different batches of mini-protein binders. The mini-proteins (sizes 5.5-14.0 kDa) were evaluated based on overall expression success rate and purity (average purity for all batches was >90%). a) Antagonist mini-protein binders designed using BindCraft, demonstrating an overall expression success rate of 93%. b) Antagonist mini-protein binders designed using RFdiffusion, demonstrating an overall expression success rate of 82%. c) Agonist mini-protein binders designed using BindCraft, demonstrating an overall expression success rate of 85%. The protein ladder used was the Seeblue Plus2 Pre-stained Standard.

**Figure S3:**
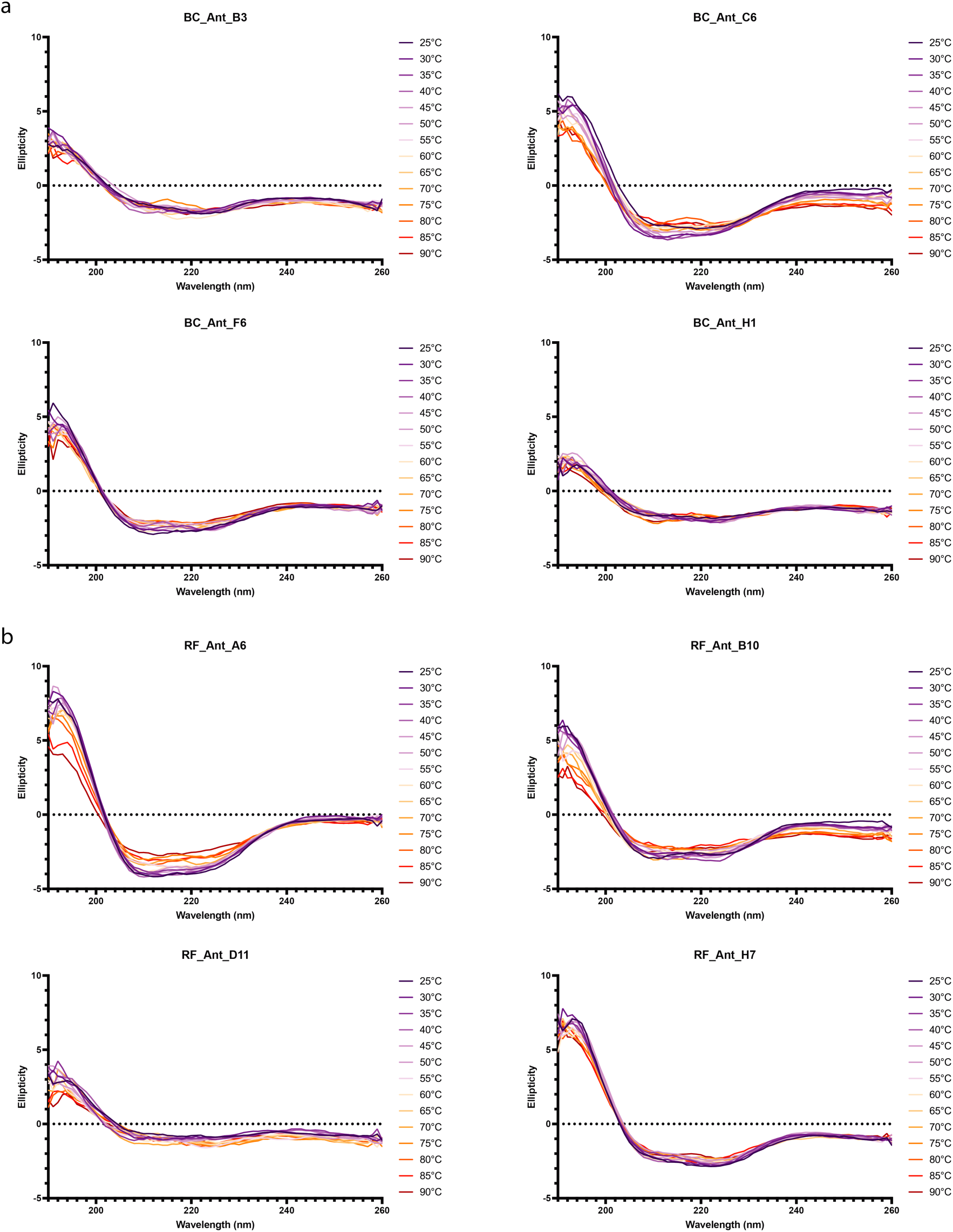
Circular dichroism spectra of antagonist mini-protein binders between 25 °C and 90 °C. a) Antagonist mini-proteins binders designed using BindCraft. **b)** Antagonist mini-protein binders using RFdiffusion. Ellipticity describes the differential absorption of left- and right-handed circularly polarised light by chiral molecules.

**Figure S4:**
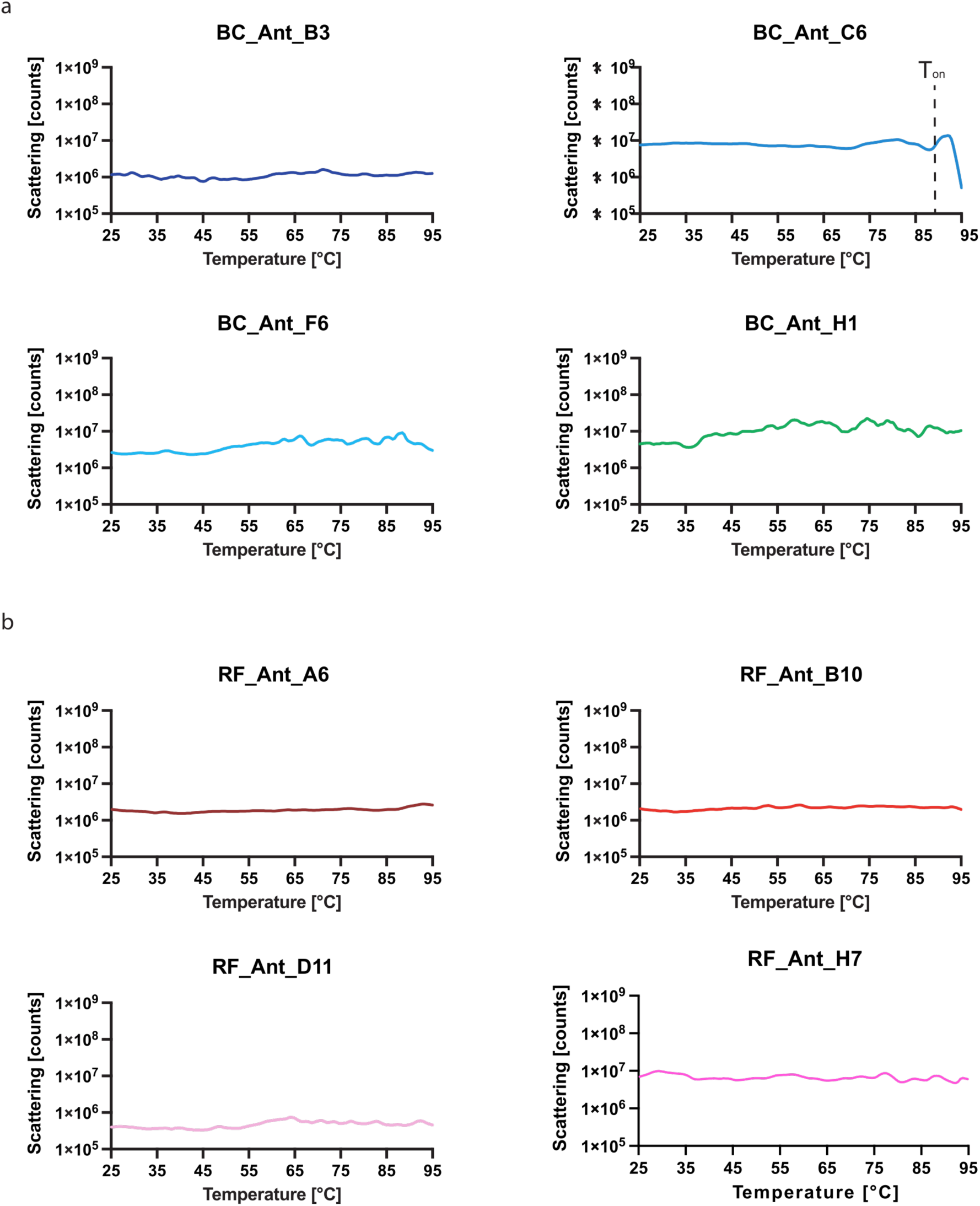
Temperature-dependent light scattering and aggregation stability of antagonist mini-protein binders. Backscattering of BindCraft **(a)** and RFdiffusion **(b)** binders monitored on a NanoTemper Prometheus instrument at 405nm over a temperature range of 25–95 °C. Dashed line labelled (T_on_) marks aggregation onset temperature.

**Figure S5:**
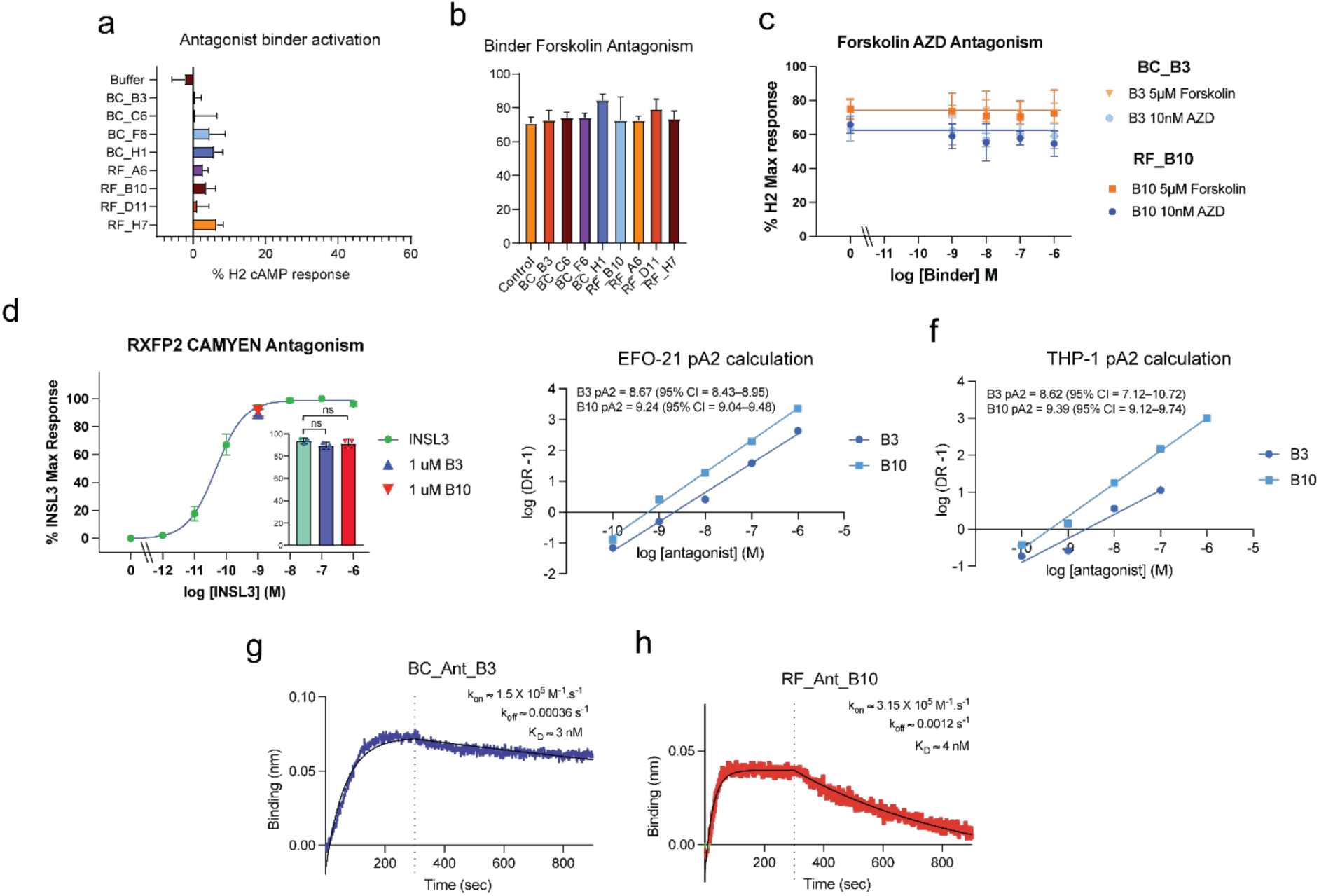
Specificity controls and binding kinetics of RXFP1 mini-protein antagonists. a) Assessment of intrins_ic_ agonist activity of selected antagonist mini-proteins in the HEK-RXFP1 CRE reporter assay. None of the antagonist binders elicited significant RXFP1 activation in the absence of H2 relaxin, indicating a lack of basal agonist activity. **b)** Effect of antagonist binders on forskolin-stimulated cAMP production in the same assay. Forskolin responses were unaffected by antagonist treatment, demonstrating that the observed inhibition of RXFP1 signalling is not due to non-specific suppression of adenylyl cyclase activity or downstream cAMP signalling. **c)** Titration of BC_Ant_B3 and RF_Ant_B10 in the presence of forskolin or the small-molecule RXFP1 agonist AZD5462. Neither antagonist altered RXFP1 activation by these compounds across the concentration range tested, consistent with selective antagonism at the extracellular domain rather than interference with transmembrane-domain-mediated activation. **d)** Activation of cAMP signalling by INSL3 in HEK293 cells expressing RXFP2, assessed via CAMYEN BRET, is not affected by 1 μM BC_Ant_B3 or RF_Ant_B10, indicating that these mini-binders do not antagonise RXFP2. **e-f)** pA_2_ calculations plots derived from equiactive dose responses (DR) of BC_Ant_B3 and RF_Ant_B10 RXFP1 antagonism curves in EFO-21 (at EC_30_) (e) and THP-1 (at EC_15_) (f) cell lines. **g,h)** Biolayer interferometry (BLI) sensorgrams showing binding kinetics of BC_Ant_B3 (g) and RF_Ant_B10 (h) to the purified RXFP1 ECD. Experimental traces are shown with fitted association–dissociation models. Derived kinetic parameters (k_on_, k_off_) and equilibrium dissociation constants (K_D) are indicated. BC_Ant_B3 displays a slower dissociation rate relative to RF_Ant_B10, consistent with its more pronounced reduction of maximal signalling observed in functional assays. Data represent mean ±S.E.M. of replicate measurements from independent experiments.

**Figure S6:**
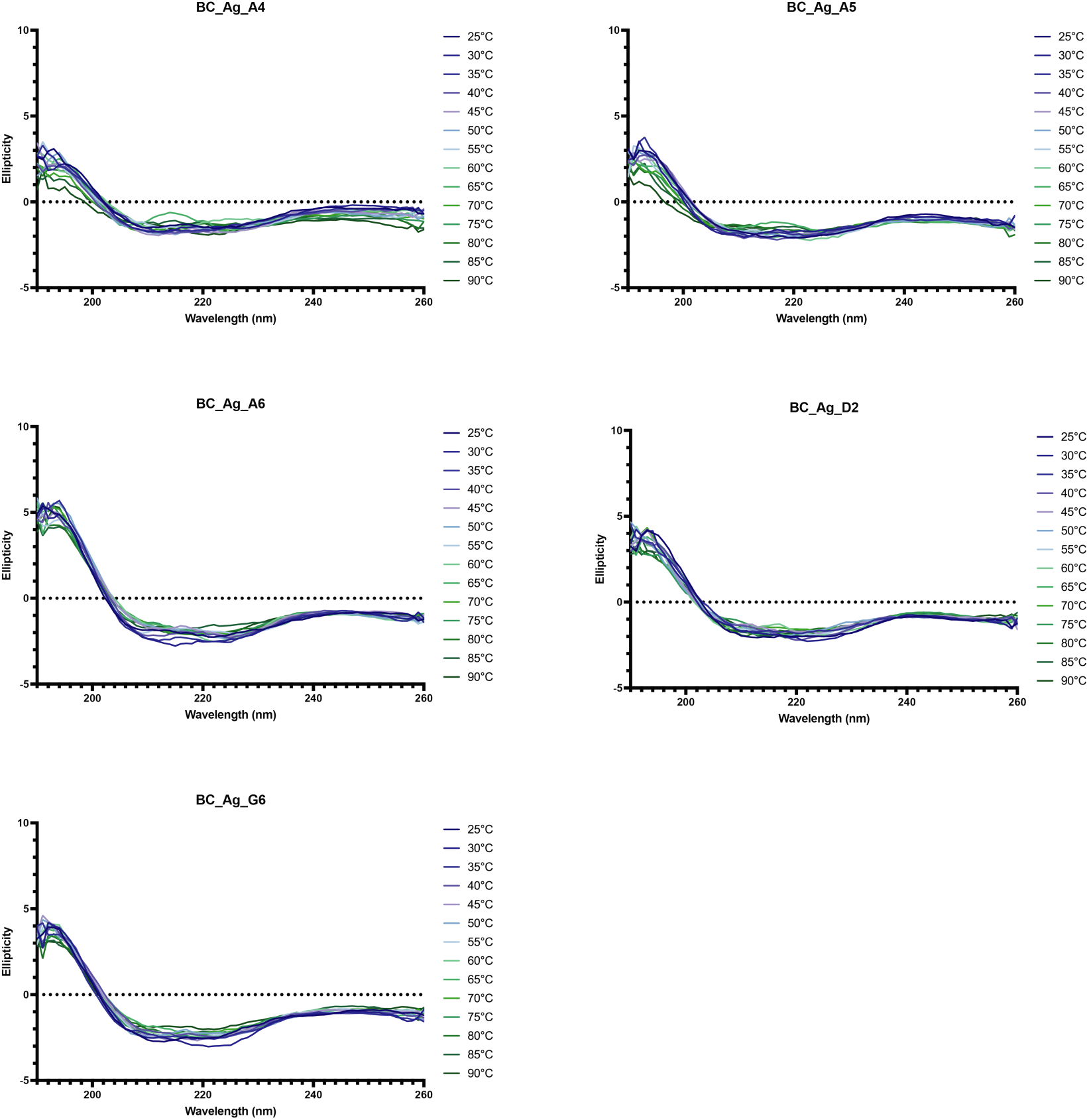
**Circular dichroism spectra of agonist mini-protein binders designed using BindCraft, recorded between 25 °C and 90 °C**. Ellipticity describes the differential absorption of left- and right-handed circularly polarised light by chiral molecules.

**Figure S7:**
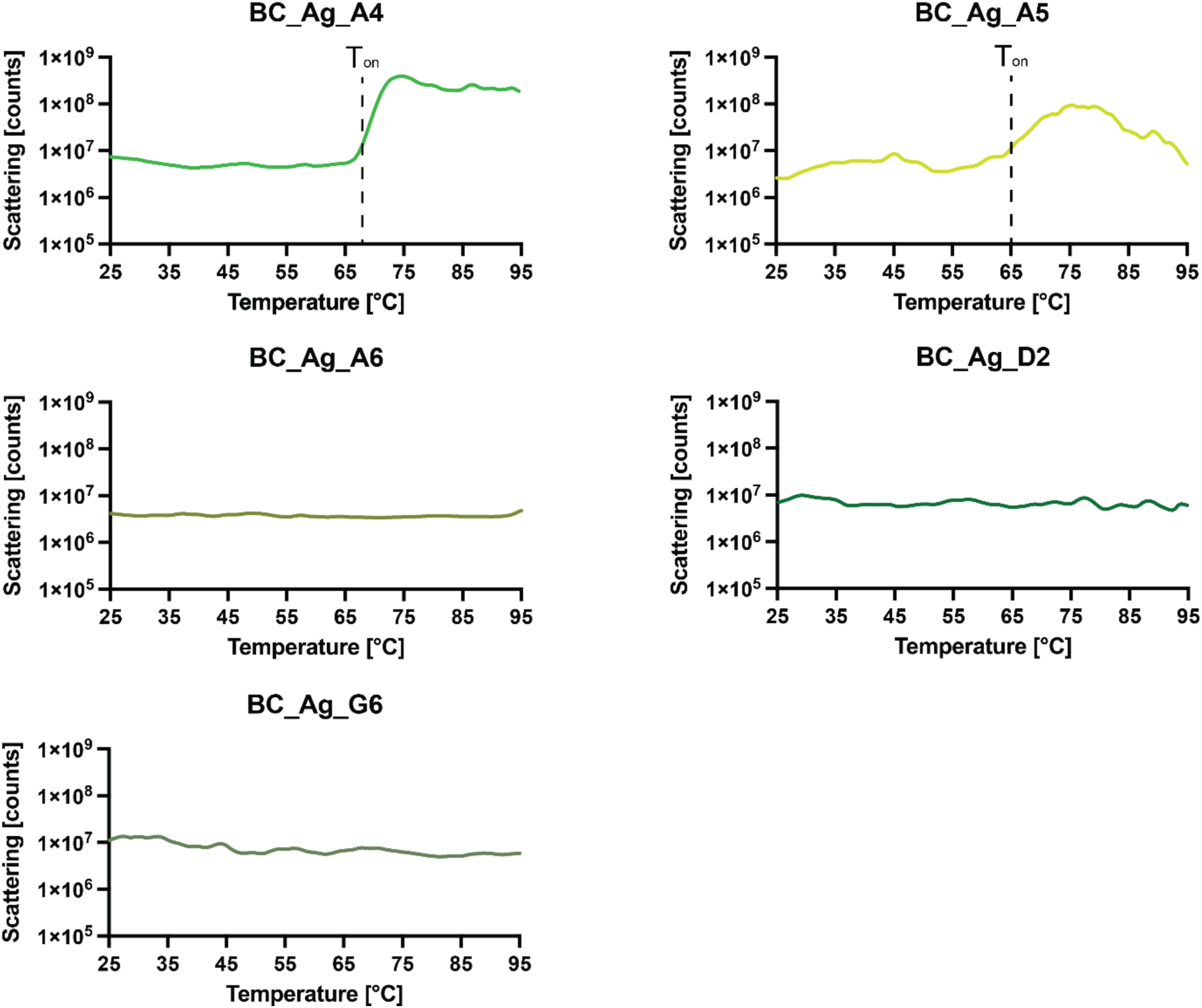
Temperature-dependent light scattering and aggregation stability of agonist mini-protein binders. Backscattering of BindCraft binders monitored on a NanoTemper Prometheus instrument at 405nm over a temperature range of 25–95 °C. Dashed lines (T_on_) mark aggregation onset temperature.

**Figure S8:**
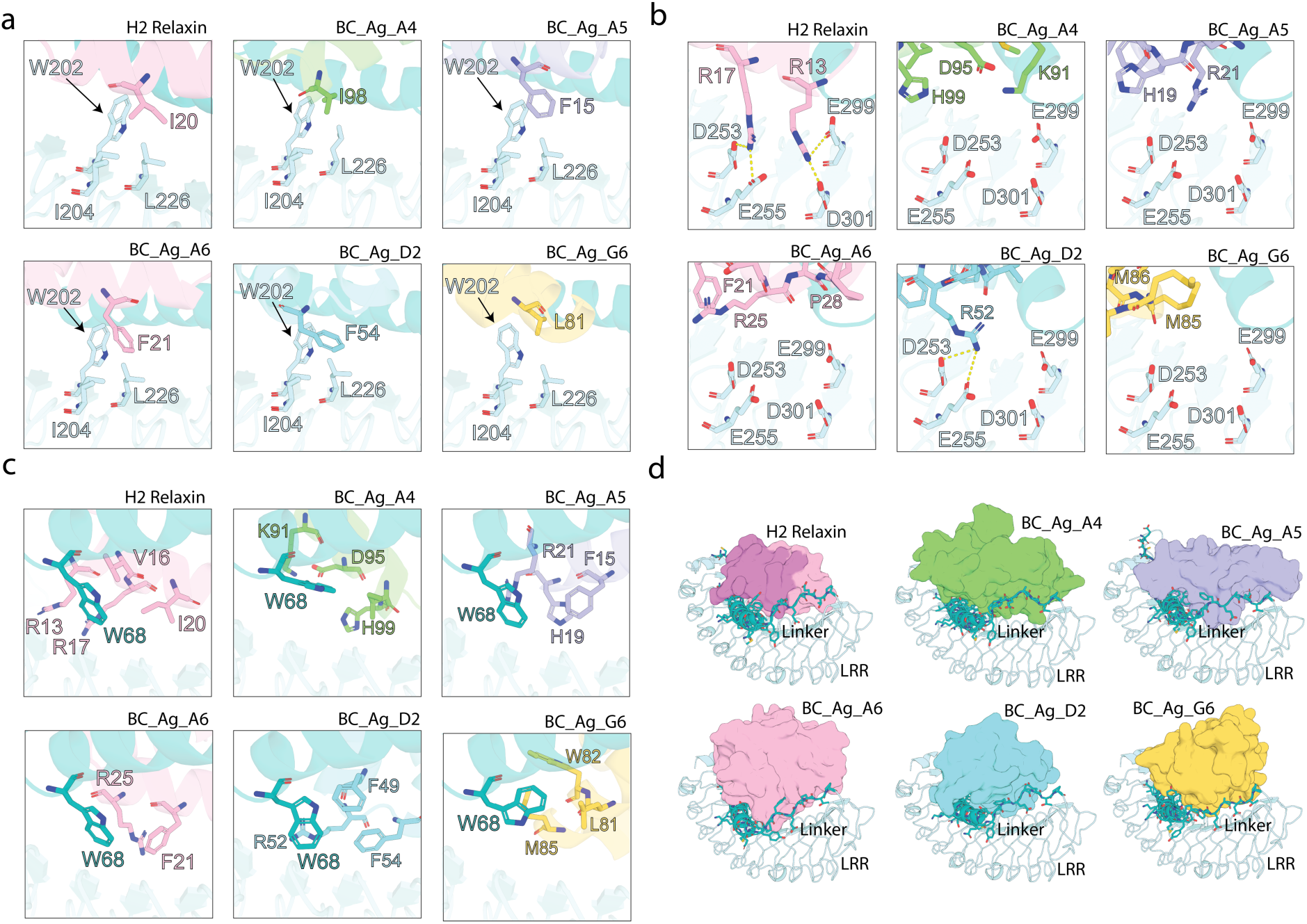
Structural comparison of relaxin and selected agonist mini-binder engagement of the RXFP1 LRR domain and linker. a) Hydrophobic interaction shared across selected agonists mimics the relaxin B-chain I20 interaction with RXFP1 W202, I204, and L226, reproduced by distinct binder residues. **b)** Key salt bridges between relaxin residues R13 and R17 and RXFP1 residues D253/E255 and E299/D301 are not conserved across the mini-binders, with the notable exception of D2 residue R52 forming a salt bridge with RXFP1 D253/E255. **c)** Convergent engagement of the RXFP1 linker residue W68 by all agonist binders, achieved through distinct interaction modes including hydrophobic packing and aromatic stacking; relaxin engages the same region using residues from the B-chain **d)** Overall binding modes of H2 relaxin and each agonist mini-binder on RXFP1, highlighting that all agonists adopt distinct folds and topologies yet converge on a similar binding surface spanning the LRR domain and the linker. Across the mini-binders, extensive contacts are formed with the RXFP1 linker region.

**Figure S9:**
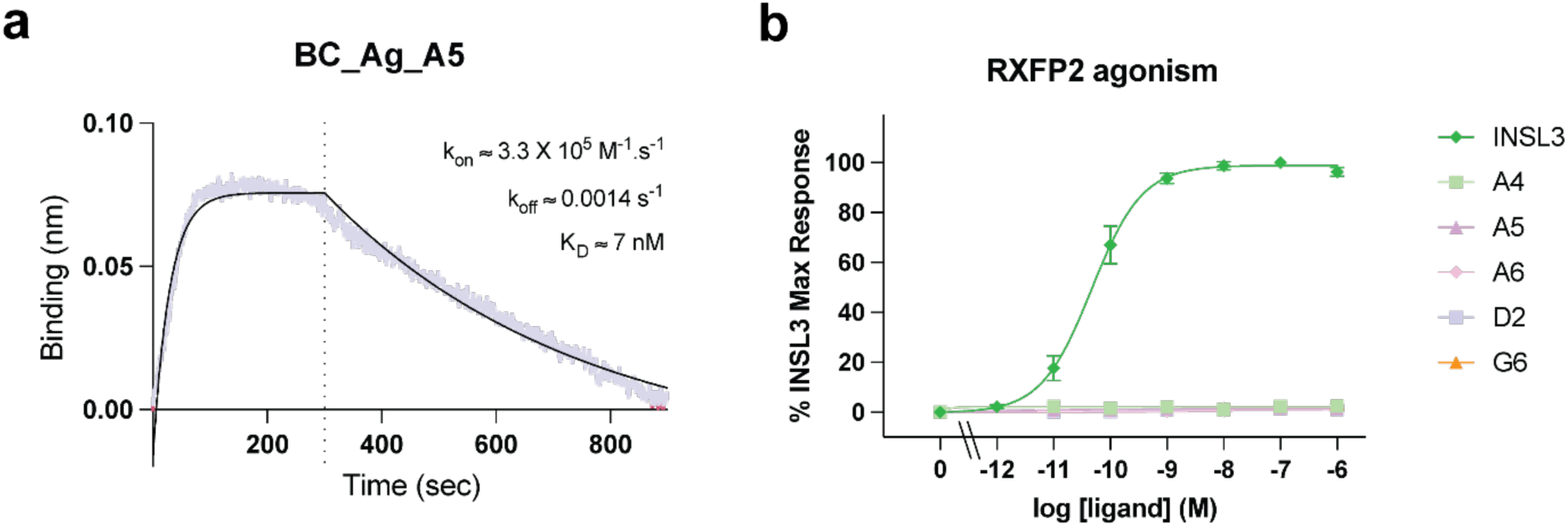
Binding affinity and specificity of RXFP1 mini-protein agonists. a) Biolayer interferometry (BLI) sensorgram showing binding kinetics of BC_Ag_A5 to the purified RXFP1 ECD. Experimental traces are shown with fitted association–dissociation models. Derived kinetic parameters (k_on_, k_off_) and equilibrium dissociation constants (K_D_) are indicated. **b)** Activation of cAMP signalling by INSL3 in HEK293 cells expressing RXFP2, assessed via CAMYEN BRET, and lack of activation by RXFP1 mini-protein agonists at 1 µM, indicating that de novo-designed agonists are highly specific for RXFP1.

## References

1 Chen, T.-Y. et al. The relaxin family peptide receptor 1 (RXFP1): An emerging player in human health and disease. Molecular Genetics & Genomic Medicine 8, e1194 (2020). 10.1002/mgg3.1194

2 Sarwar, M., Du, X.-J., Dschietzig, T. B. & Summers, R. J. The actions of relaxin on the human cardiovascular system. British Journal of Pharmacology 174, 933–949 (2017). 10.1111/bph.13523

3 Samuel, C. S. et al. Anti-fibrotic actions of relaxin. British Journal of Pharmacology 174, 962–976 (2017). 10.1111/bph.13529

4 Du, X.-J., Bathgate, R. A. D., Samuel, C. S., Dart, A. M. & Summers, R. J. Cardiovascular effects of relaxin: from basic science to clinical therapy. Nature Reviews Cardiology 7, 48–58 (2010). 10.1038/nrcardio.2009.198

5 Sethi, A. et al. Structural Insights into the Unique Modes of Relaxin-Binding and Tethered-Agonist Mediated Activation of RXFP1 and RXFP2. Journal of Molecular Biology 433, 167217 (2021). 10.1016/j.jmb.2021.167217

6 Petrie, E. J., Lagaida, S., Sethi, A., Bathgate, R. A. & Gooley, P. R. In a class of their own–RXFP1 and RXFP2 are unique members of the LGR family. Frontiers in endocrinology 6, 137 (2015).

7 Hsu, S. Y. et al. Activation of Orphan Receptors by the Hormone Relaxin. Science 2G5, 671–674 (2002). doi:10.1126/science.1065654

8 Bathgate, R. et al. Relaxin family peptides and their receptors. Physiological reviews **G3**, 405–480 (2013).

9 Bathgate, R. A. D. et al. The relaxin receptor as a therapeutic target - perspectives from evolution and drug targeting. Pharmacol Ther 187, 114–132 (2018). 10.1016/j.pharmthera.2018.02.008

10 Bani, D. Recombinant human H2 relaxin (serelaxin) as a cardiovascular drug: aiming at the right target. Drug Discovery Today 25, 1239–1244 (2020). 10.1016/j.drudis.2020.04.014

11 Borlaug, B. A. et al. Effects of volenrelaxin in worsening heart failure with preserved ejection fraction: a phase 2 randomized trial. Nature Medicine 31, 3853–3861 (2025). 10.1038/s41591-025-03939-6

12 Erlandson, S. C. et al. Engineering and Characterization of a Long-Half-Life Relaxin Receptor RXFP1 Agonist. Molecular Pharmaceutics 21, 4441–4449 (2024). 10.1021/acs.molpharmaceut.4c00368

13 Kanai, A. J., Konieczko, E. M., Bennett, R. G., Samuel, C. S. & Royce, S. G. Relaxin and fibrosis: Emerging targets, challenges, and future directions. Molecular and Cellular Endocrinology 487, 66–74 (2019). 10.1016/j.mce.2019.02.005

14 Ufnal, M. et al. Relaxin mimetic in pulmonary hypertension associated with left heart disease: Design and rationale of Re-PHIRE. ESC Heart Failure 12, 1956–1964 (2025). 10.1002/ehf2.15203

15 Papworth, M. et al. A novel long-acting relaxin-2 fusion, AZD3427, improves cardiac performance in non-human primates with cardiac dysfunction. Cardiovascular Research 121, 871–881 (2025).

16 Hossain, M. A. et al. Development of novel high-affinity antagonists for the relaxin family peptide receptor 1. ACS Pharmacology & Translational Science 6, 842–853 (2023).

17 Ilyas, S. I. & Gores, G. J. The Two Faces of Relaxin in Cancer: Antitumor or Protumor? Hepatology 71, 1117–1119 (2020). 10.1002/hep.30998

18 Burston, H. E. et al. Inhibition of relaxin autocrine signaling confers therapeutic vulnerability in ovarian cancer. The Journal of Clinical Investigation 131 (2021). 10.1172/JCI142677

19 Almeida-Pinto, N., Dschietzig, T. B., Bras-Silva, C. & Adão, R. Cardiovascular effects of relaxin-2: therapeutic potential and future perspectives. Clinical Research in Cardiology 113, 1137–1150 (2024).

20 Tham, L. S. et al. Volenrelaxin (LY3540378) increases renal plasma flow: a randomized Phase 1 trial. Nephrology Dialysis Transplantation 40, 109–122 (2025).

21 Tan, J. et al. Expression of RXFP1 Is Decreased in Idiopathic Pulmonary Fibrosis. Implications for Relaxin-based Therapies. Am J Respir Crit Care Med 194, 1392–1402 (2016). 10.1164/rccm.201509-1865OC

22 Erlandson, S. C. et al. The relaxin receptor RXFP1 signals through a mechanism of autoinhibition. Nature Chemical Biology **19**, 1013–1021 (2023). 10.1038/s41589-023-01321-6

23. Fox, D. R., Taveneau, C., Clement, J., Grinter, R. & Knott, G. J. Code to complex: AI-driven <em>de novo</em> binder design. *Structure* 33, 1631–1642 (2025). 10.1016/j.str.2025.08.007

24 Fox, D. R. et al. Inhibiting heme piracy by pathogenic Escherichia coli using de novo-designed proteins. Nature Communications 16, 6066 (2025). 10.1038/s41467-025-60612-9

25 Jumper, J. et al. Highly accurate protein structure prediction with AlphaFold. nature 5G6, 583–589 (2021).

26 Evans, R. et al. Protein complex prediction with AlphaFold-Multimer. bioRxiv, 2021.2010.2004.463034 (2022). 10.1101/2021.10.04.463034

27 Abramson, J. et al. Accurate structure prediction of biomolecular interactions with AlphaFold 3. Nature 630, 493–500 (2024).

28 Lyu, J. et al. AlphaFold2 structures guide prospective ligand discovery. Science 384, eadn6354 (2024). doi:10.1126/science.adn6354

29 Sethi, A. et al. The complex binding mode of the peptide hormone H2 relaxin to its receptor RXFP1. Nature Communications 7, 11344 (2016). 10.1038/ncomms11344

30 Hoare, B. L. et al. Multi-Component Mechanism of H2 Relaxin Binding to RXFP1 through NanoBRET Kinetic Analysis. iScience 11, 93–113 (2019). 10.1016/j.isci.2018.12.004

31 Watson, J. L. et al. De novo design of protein structure and function with RFdiffusion. Nature 620, 1089–1100 (2023).

32 Pacesa, M. et al. One-shot design of functional protein binders with BindCraft. Nature 646, 483–492 (2025).

33 Dauparas, J. et al. Robust deep learning–based protein sequence design using ProteinMPNN. Science 378, 49–56 (2022).

34 Bennett, N. R. et al. Improving de novo protein binder design with deep learning. Nature Communications 14, 2625 (2023).

35 Büllesbach, E. E. & Schwabe, C. The trap-like relaxin-binding site of the leucine-rich G-protein-coupled receptor 7. Journal of Biological Chemistry 280, 14051–14056 (2005).

36 Rosengren, K. J. et al. Solution structure and novel insights into the determinants of the receptor specificity of human relaxin-3. J Biol Chem 281, 5845–5851 (2006). 10.1074/jbc.M511210200

37 Eigenbrot, C. et al. X-ray structure of human relaxin at 1· 5Å: Comparison to insulin and implications for receptor binding determinants. Journal of molecular biology 221, 15–21 (1991).

38 Yan, Y. et al. Identification of the N-linked glycosylation sites of the human relaxin receptor and effect of glycosylation on receptor function. Biochemistry 47, 6953–6968 (2008). 10.1021/bi800535b

39 Hu, X. et al. Structural Insights into the Activation of Human Relaxin Family Peptide Receptor 1 by Small-Molecule Agonists. Biochemistry 55, 1772–1783 (2016). 10.1021/acs.biochem.5b01195

40 Granberg, K. L. et al. Discovery of Clinical Candidate AZD5462, a Selective Oral Allosteric RXFP1 Agonist for Treatment of Heart Failure. Journal of Medicinal Chemistry 67, 4419–4441 (2024). 10.1021/acs.jmedchem.3c02184

41 Ghandi, M. et al. Next-generation characterization of the Cancer Cell Line Encyclopedia. Nature **569**, 503–508 (2019). 10.1038/s41586-019-1186-3

42 Kenakin, T., Jenkinson, S. & Watson, C. Determining the potency and molecular mechanism of action of insurmountable antagonists. The Journal of pharmacology and experimental therapeutics **319**, 710–723 (2006).

43 Broszkiewicz, W. & Domińska, K. The role of relaxins in blood cell modulation: interactions with relaxin family peptide receptor 1 (RXFP1) and glucocorticoid receptor (GR). Clinical Science **139**, 1431–1450 (2025). 10.1042/cs20256619

44 Fleit, H. B. & Kobasiuk, C. D. The human monocyte-like cell line THP-1 expresses FcγRI and FCγRII. Journal of leukocyte biology **49**, 556–565 (1991).

45 Scott, D. J., Rosengren, K. J. & Bathgate, R. A. The different ligand-binding modes of relaxin family peptide receptors RXFP1 and RXFP2. Molecular Endocrinology 26, 1896–1906 (2012).

46 Vauquelin, G. & Charlton, S. J. Long-lasting target binding and rebinding as mechanisms to prolong in vivo drug action. British journal of pharmacology 161, 488–508 (2010).

47 Hellmann, J. et al. Structure-based development of a subtype-selective orexin 1 receptor antagonist. Proceedings of the National Academy of Sciences 117, 18059–18067 (2020).

48 Riddy, D. M. et al. Label-free kinetics: exploiting functional hemi-equilibrium to derive rate constants for muscarinic receptor antagonists. Molecular Pharmacology 88, 779–790 (2015).

49 Kern, A. & Bryant-Greenwood, G. D. Characterization of relaxin receptor (RXFP1) desensitization and internalization in primary human decidual cells and RXFP1-transfected HEK293 cells. Endocrinology 150, 2419–2428 (2009).

50 Elfmann, C. & Stülke, J. PAE viewer: a webserver for the interactive visualization of the predicted aligned error for multimer structure predictions and crosslinks. Nucleic Acids Res 51, W404–w410 (2023). 10.1093/nar/gkad350

51 Gasteiger, E. et al. in The proteomics protocols handbook 571–607 (Springer, 2005).

52 Halls, M. L., Bathgate, R. A. & Summers, R. J. Comparison of signaling pathways activated by the relaxin family peptide receptors, RXFP1 and RXFP2, using reporter genes. The Journal of pharmacology and experimental therapeutics 320, 281–290 (2007).

53 Hossain, M. A. et al. The A-chain of human relaxin family peptides has distinct roles in the binding and activation of the different relaxin family peptide receptors. Journal of Biological Chemistry 283, 17287–17297 (2008).

54 Valkovic, A. L. et al. Real-time examination of cAMP activity at relaxin family peptide receptors using a BRET-based biosensor. Pharmacol Res Perspect 6, e00432 (2018). 10.1002/prp2.432

55 Jiang, L. I. et al. Use of a cAMP BRET sensor to characterize a novel regulation of cAMP by the sphingosine 1-phosphate/G13 pathway. J Biol Chem 282, 10576–10584 (2007). 10.1074/jbc.M609695200

56 Mönnich, D. et al. Activation of multiple G protein pathways to characterize the five dopamine receptor subtypes using bioluminescence technology. ACS Pharmacology & Translational Science 7, 834–854 (2024).

57 Hossain, M. A. et al. A single-chain derivative of the relaxin hormone is a functionally selective agonist of the G protein-coupled receptor, RXFP1. Chemical Science 7, 3805–3819 (2016). 10.1039/C5SC04754D

58 Götzke, H. et al. The ALFA-tag is a highly versatile tool for nanobody-based bioscience applications. Nature Communications 10, 4403 (2019). 10.1038/s41467-019-12301-7

59 Kariuki, C. K. & Magez, S. Improving the yield of recalcitrant Nanobodies® by simple modifications to the standard protocol. Protein Expression and Purification 185, 105906 (2021).

